# An assessment of environmental metabarcoding protocols aiming at favouring contemporary biodiversity in inventories of deep-sea communities

**DOI:** 10.1101/836080

**Authors:** Miriam I. Brandt, Blandine Trouche, Nicolas Henry, Cathy Liautard-Haag, Lois Maignien, Colomban de Vargas, Patrick Wincker, Julie Poulain, Daniela Zeppilli, Sophie Arnaud-Haond

## Abstract

1. The abyssal seafloor covers more than 50% of planet Earth and is a large reservoir of still mostly undescribed biodiversity. It is increasingly under target of resource-extraction industries although being drastically understudied. In such remote and hard-to-access ecosystems, environmental DNA (eDNA) metabarcoding is a useful and efficient tool for studying biodiversity and implementing environmental impact assessments. Yet, eDNA analysis outcomes may be biased towards describing past rather than present communities as sediments contain both contemporary and ancient DNA.
2. Using commercially available kits, we investigated the impacts of five molecular processing methods on DNA metabarcoding biodiversity inventories targeting prokaryotes (16S-V4V5), unicellular eukaryotes (18S-V4), and metazoans (18S-V1, COI). As the size distribution of ancient DNA is skewed towards small fragments, we evaluated the effect of removing short DNA fragments via size-selection and ethanol reconcentration using DNA extracted from 10 g of sediment at five deep-sea sites. We also compare communities revealed by DNA and RNA co-extracted from 2 g of sediment at the same sites.
3. Results show that removing short DNA fragments does not affect alpha and beta diversity estimates in any of the biological compartments investigated. Results also confirm doubts regarding the possibility to better describe *live* communities using environmental RNA (eRNA). With ribosomal loci, RNA, while resolving similar spatial patterns than co-extracted DNA, resulted in significantly higher richness estimates, supporting hypotheses of increased persistence of ribosomal RNA (rRNA) in the environment and unmeasured bias due to over-abundance of rRNA and RNA release. With the mitochondrial locus, RNA detected lower metazoan richness and resolved less spatial patterns than co-extracted DNA, reflecting high messenger RNA lability. Results also highlight the importance of using large amounts of sediment (≥10 g) for accurately surveying eukaryotic diversity.
4. We conclude that DNA should be favoured over RNA for logistically realistic, repeatable, and reliable surveys, and confirm that large sediment samples (≥10 g) deliver more complete and accurate assessments of benthic eukaryotic biodiversity and that increasing the number of biological rather than technical replicates is important to infer robust ecological patterns.

## 1. Introduction

Environmental DNA (eDNA) metabarcoding is an increasingly used tool for biodiversity inventories and ecological surveys. Using high-throughput sequencing (HTS) and bioinformatic processing, it allows the detection or the inventory of target organisms using their DNA directly extracted from soil, water, or air samples (Taberlet, Coissac, Hajibabaei, & Rieseberg, 2012). As it does not require specimen isolation, it represents a practical and efficient tool in large and hard-to-access ecosystems, such as the marine realm. Besides allowing studying various biological compartments simultaneously, metabarcoding is also very effective for detecting diversity of small organisms (micro-organisms, meiofauna) largely disregarded in visual biodiversity inventories due to the difficulty of their identification based on morphological features (Carugati, Corinaldesi, Dell’Anno, & Danovaro, 2015).

The deep sea, covering more than 50% of Planet Earth, remains critically understudied, despite being increasingly impacted by anthropogenic activities and targeted by resource-extraction industries (Ramirez-Llodra et al., 2011). The abyssal seafloor is mostly composed of sedimentary habitats containing high numbers of small (< 1 mm) organisms, and characterized by high local and regional diversity (Grassle & Maciolek, 1992; Smith & Snelgrove, 2002). Given the increased time-efficiency offered by eDNA metabarcoding and its wide taxonomic applicability, this tool is a good candidate for large-scale biodiversity surveys and Environmental Impact Assessments (EIAs) in the deep-sea biome.

eDNA is a complex mixture of genomic DNA present in living cells, extra-organismal DNA, and extracellular DNA originating from the degradation of organic material and biological secretions (Torti, Lever, & Jørgensen, 2015). Extracellular DNA has been shown to be very abundant in marine sediments, representing 50-90% of the total DNA pool (Corinaldesi, Tangherlini, Manea, & Dell’Anno, 2018; Dell’Anno & Danovaro, 2005). However, this extracellular DNA compartment may not only contain DNA from contemporary communities. Indeed, nucleic acids can persist in marine sediments as their degradation rate decreases due to adsorption onto the sediment matrix (Corinaldesi, Beolchini, & Dell’Anno, 2008; Torti et al., 2015). Low temperatures, high salt concentrations, and the absence of UV light are additional factors enhancing long-term archiving of DNA in deep-sea sediments (Nagler, Insam, Pietramellara, & Ascher-Jenull, 2018; Torti et al., 2015). Decreased rates of abiotic DNA decay can thus allow DNA persistence over millennial timescales. Indeed, up to 125,000-year-old ancient DNA (aDNA) has been reported in oxic and anoxic marine sediments at various depths (Boere, Rijpstra, De Lange, Sinninghe Damsté, & Coolen, 2011; Coolen et al., 2013; Lejzerowicz, Esling, et al., 2013). As extracellular DNA fragment size depends on its state of degradation (Nagler et al., 2018 report overall size ranges from 80 to over 20,000 bp), aDNA fragments have generally been reported to be <1,000 bp long (Boere et al., 2011; Coolen et al., 2013; Lejzerowicz, Esling, et al., 2013; Lennon, Muscarella, Placella, & Lehmkuhl, 2018). Restricting molecular biodiversity assessments to large DNA fragments may thus allow avoiding the bias of aDNA in biodiversity assessments aiming at describing contemporary communities using eDNA metabarcoding.

Environmental RNA (eRNA) has been viewed as a way to avoid the problem of aDNA in eDNA biodiversity inventories because RNA is only produced by living organisms and quickly degrades when released in the environment, due to spontaneous hydrolysis and the abundance of RNases (Torti et al., 2015). Few studies have investigated this in the deep-sea, with contrasting results. Investigating foraminiferal assemblages, Lejzerowicz, Voltsky, & Pawlowski (2013) found similar taxonomic compositions with DNA and RNA, although highlighting that RNA is more appropriate for targeting the active community component. Contrastingly, Guardiola et al. (2016) detected marked differences between RNA and DNA inventories for most eukaryotic groups, but found that both biomolecules detected similar patterns of ecological differentiation, concluding that “dead” DNA did not blur patterns of community structure. Laroche and coworkers (2018, 2017) found stronger responses to environmental impact in alpha diversity measured with eRNA, while eDNA was better at detecting effects on community composition. Finally, long-term archived and even fossil RNA were also reported in sediment and soil (Cristescu, 2019; Orsi, Biddle, & Edgcomb, 2013), casting doubts as to its advantage over DNA to inventory contemporary biodiversity.

The design of a sound environmental metabarcoding protocol to inventory biodiversity on the deep seafloor relies on a better understanding of the potential influence of aDNA on the different taxonomic compartments targeted. Using commercially available kits based on 2 g and 10 g of sediment, we studied samples from five deep-sea sites encompassing three different habitats and spanning wide geographic ranges, in order to select an optimal protocol to survey contemporary benthic deep-sea communities spanning the tree of life. We analyse eDNA and eRNA extracts via metabarcoding, targeting the 16S ribosomal RNA (rRNA) barcode (Parada, Needham, & Fuhrman, 2016) for prokaryotes, the 18S-V4 rRNA barcode region for micro-eukaryotes (Stoeck et al., 2010), and the 18S-V1V2 rRNA and COI barcode markers (Leray et al., 2013; Sinniger et al., 2016) for metazoans.

Our objectives were threefold:

1. Evaluate the effect of removing short DNA fragments from DNA extracts obtained using a 10 g extraction kit;
2. Compare eDNA and eRNA inventories resulting from the same samples via a 2 g joint extraction kit,
3. Assess the aforementioned kits in terms of repeatability and suitability for different taxonomic compartments.

## 2. Materials and methods

### 2.1. Collection of samples

Sediment cores were collected from five deep-sea sites from various habitats (mud volcano, seamounts, and an area close to hydrothermal vents, Table S 1). Triplicate tube cores were collected with a multicorer or with a remotely operated vehicle at each sampling site. The sediment cores were sliced into layers, which were transferred into zip-lock bags, homogenised, and frozen at −80°C on board before being shipped on dry ice to the laboratory. The first layer (0-1 cm) was used for the present analysis. In each sampling series, an empty bag was kept as a field control processed through DNA extraction and sequencing.

### 2.2. Nucleic acid extractions and molecular treatments

#### eDNA from 10 g of sediment

DNA extractions were performed using ∼10 g of sediment with the PowerMax Soil DNA Isolation Kit (MO BIO Laboratories, Inc.; Qiagen, Hilden, Germany). To increase the DNA yield, the elution buffer was left on the spin filter membrane for 10 min at room temperature before centrifugation. For field controls, the first solution of the kit was poured into the control zip lock, before following the usual extraction steps. DNA extracts were stored at −80°C.

#### Size-selection of eDNA extracts

Size-selection of total eDNA extracted as detailed above from 10 g of sediment was carried out to remove small DNA fragments. NucleoMag NGS Clean-up and Size Select beads (Macherey-Nagel, Düren, Germany) were used at a ratio of 0.5X for removing DNA fragments < 1,000 bp from 500 µL of extracted eDNA. The target fragments were eluted from the beads with 100 µL elution buffer, and successful size-selection verified by electrophoresis on an Agilent TapeStation using the Genomic DNA High ScreenTape kit (Agilent Technologies, Santa Clara, CA, USA).

#### Ethanol reconcentration of eDNA extracts

A 3.5 mL aliquot of eDNA extracted from 10 g of sediment was reconcentrated with 7 mL of 96% ethanol and 200 µl of 5M NaCl, according to the guidelines in the *Hints and Troubleshooting Guide* of the PowerMax Soil DNA Isolation Kit. As this protocol does not include any incubation time, it favours large DNA fragments. The DNA pellet was washed with 1 mL 70% ethanol, centrifuged again for 15 min at 2500 × g, and air-dried before being resuspended in 450 µL elution buffer.

#### Joint environmental DNA/RNA from 2 g of sediment

Joint RNA/DNA extractions were performed with the RNA PowerSoil Total RNA Isolation Kit combined with the RNeasy PowerSoil DNA elution kit (MO BIO Laboratories, Inc.; Qiagen, Hilden, Germany). About 5 g of sediment were used, following the manufacturer’s suggestions for wet sediments. Extraction controls were performed alongside sample extractions. The RNA pellet was resuspended in 60 uL of RNase/DNase-free water. Extracted RNA was then transcribed to first-strand complementary DNA (cDNA) using the iScript Select cDNA synthesis kit (Bio-Rad laboratories, CA, USA) with its proprietary random primer mix. Quality control 16S-V4V5, 18S-V1, and COI PCRs were performed on the RNA extracts to test for potential DNA contamination.

### 2.3. PCR amplification and sequencing

Four primer pairs were used to amplify one mitochondrial and three rRNA barcode loci targeting metazoans (COI, 18S-V1), micro-eukaryotes (18S-V4) and prokaryotes (16S-V4V5, Table S 2). Two metazoan mock communities (detailed in Brandt et al., 2019) were included for 18S-V1 and COI. For each sample and marker, triplicate amplicon libraries (see Supporting Information for amplification details) were prepared by ligation of Illumina adapters on 100 ng of amplicons following the Kapa Hifi HotStart NGS library Amplification kit (Kapa Biosystems, Wilmington, MA, USA). After quantification and quality control, library concentrations were normalized to 10 nM, and 8–9 pM of each library containing a 20% PhiX spike-in were sequenced on a HiSeq2500 (System User Guide Part # 15035786) instruments in a 250 bp paired-end mode.

### 2.4. Bioinformatic analyses

All bioinformatic analyses were performed using a Unix shell script (Brandt et al., 2019), available on Gitlab (https://gitlab.ifremer.fr/abyss-project/), on a home-based cluster (DATARMOR, Ifremer). Pairs of Illumina reads were corrected with DADA2 v.1.10 (Callahan et al., 2016) following the online tutorial for paired-end data (https://benjjneb.github.io/dada2/tutorial.html). The details of the pipeline, along with specific parameters used for all metabarcoding markers, are given in Table S 3 and in Brandt et al. (2019).

Micro-eukaryote and prokaryote diversity was evaluated with Amplicon Sequence Variants (ASVs), while metazoan data was further clustered into OTUs with the FROGS pipeline (Escudié et al., 2018). ASVs and OTUs were taxonomically assigned via BLAST+ (v2.6.0) based on minimum similarity and minimum coverage (-perc_identity 70 and – qcov_hsp 80). The Silva132 reference database was used for taxonomic assignment of rRNA marker genes (Quast et al., 2012), and MIDORI-UNIQUE (Machida, Leray, Ho, & Knowlton, 2017) was used for COI.

Molecular inventories were refined in R v.3.5.1 (R Core Team, 2018). A blank correction was made using the *decontam* package v.1.2.1 (Davis, Proctor, Holmes, Relman, & Callahan, 2018), removing all clusters that were more prevalent in negative control samples than in true or mock samples. Unassigned and non-target clusters were removed. Additionally, for metazoan loci, all clusters with a terrestrial assignment (groups known to be terrestrial-only) were removed. Samples with less than 10,000 target reads were discarded. We performed an abundance renormalization to remove spurious ASVs/OTUs due to random tag switching (Wangensteen & Turon, 2016). The COI OTU table was further curated with LULU v.0.1 (Frøslev et al., 2017) to limit the bias due to pseudogenes, using a minimum co-occurrence of 0.93 and a minimum similarity threshold of 84%.

### 2.5. Statistical analyses

Sequence tables were analysed using R with the packages phyloseq v1.22.3 (McMurdie & Holmes, 2013), following guidelines in online tutorials (http://joey711.github.io/phyloseq/tutorials-index.html), and vegan v2.5.2 (Oksanen et al., 2018). Alpha diversity between molecular processing methods was estimated with the number of observed target clusters in rarefied datasets. Cluster abundances were compared via analyses of deviances (ANODEV) on generalized linear mixed models using negative binomial distributions, as the data were overdispersed. Pairwise post-hoc comparisons were performed via Tukey HSD tests using the *emmeans* package.

Homogeneity of multivariate dispersions were evaluated with the *betapart* package v.1.5.1 (Baselga & Orme, 2012), and statistical tests performed on balanced datasets for COI as dispersions were different between 2 g and 10g datasets (Table S 4). Data were rarefied for metazoans and Hellinger-normalised for microbial data.

Differences in community compositions resulting from molecular processing were evaluated with Mantel tests (Jaccard and Bray-Curtis dissimilarities for metazoan and microbial taxa respectively; Pearson’s product–moment correlation; 1000 permutations). Permutational multivariate analysis of variance (PERMANOVA) was performed on normalised datasets to evaluate the effect of molecular processing and site on community compositions, using the function *adonis2* (vegan) with Jaccard dissimilarities (presence/absence) for metazoan and Bray-Curtis dissimilarities for prokaryotes and micro-eukaryotes. The rationale behind this choice is that metazoans are multicellular organisms of extremely varying numbers of cells, organelles, or ribosomal repeats in their genomes, and can also be detected through a diversity of remains. The number of reads can thus not be expected to reflect the abundance of detected OTUs. Significance was evaluated via marginal effects of terms, using 10,000 permutations with site as a blocking factor. Pairwise post-hoc comparisons were performed via the *pairwiseAdonis* package, with site as a blocking factor. Differences between samples were visualized via Principal Coordinates Analyses (PCoA) based on abovementioned dissimilarities. Finally, taxonomic compositions in terms of cluster and read abundance were compared between molecular processing methods on balanced datasets.

## 3. Results

### 3.1. High-throughput sequencing results

A total of 70 million 18S-V1 reads, 61 million COI reads, 30 million 18S-V4 reads, and 45 million 16S-V4V5 reads were obtained from four Illumina HiSeq runs of pooled amplicon libraries built from triplicate PCR replicates of 75 sediment samples, 2 mock communities (for 18S-V1 and COI), 3 extraction blanks, and 2-4 PCR negative controls(Table S 5). One to seven sediment samples failed amplification in each dataset. These were always coming from the same sampling sites (MDW-ST117 and MDW-ST38), and predominantly comprised RNA samples (Table S 5). After bioinformatic processing, read numbers were reduced to 44 million for 18S-V1, 45 million for COI, 16 million for 18S-V4, and 24 million for 16S-V4V5 (Table S 5). For eukaryote markers, less reads were retained in negative controls (2-64%) than in true or mock samples (49-83%), while the opposite was observed for 16S-V4V5 (62% of reads retained in control samples against 49-57% in true samples). Negative control samples (extraction and PCR blanks) contained 0.001-0.6% of total processed reads, compared to 1.3-1.5% in a true samples.

DNA extracts obtained from the joint DNA/RNA protocol based on 2 g of sediment produced less eukaryotic reads than DNA extracts from the 10 g kit, while similar yields were obtained for prokaryotes. RNA extracts produced more reads than DNA extracts with the ribosomal loci, while they produced less reads with the mitochondrial COI locus (Table S 5).

After data refining and abundance renormalisation (Wangensteen & Turon, 2016), the final datasets comprised between 8.6 and 16.2 million target reads for eukaryotes and 21.7 million prokaryote reads for 16S-V4V5. Target reads delivered 4,333 and 6,031 metazoan OTUs for COI and 18S-V1 respectively, 40,868 micro-eukaryote 18S-V4 ASVs, and 138,478 prokaryote 16S ASVs (Table S5).

### 3.2. Alpha diversity between processing methods

Rarefaction curves showed a plateau was reached for all samples, suggesting an overall sequencing depth adequate to capture the diversity present (Fig. S 1). Processing methods significantly affected the number of recovered eukaryote and prokaryote clusters, and significant variability among sites was detected for 18S-V1V2 and 18S-V4 (Table 1, Figs 1-S 2).

**Table 1.** ANODEVs and PERMANOVAs for the four studied genes. The ANODEVs were performed on mixed models with negative binomial distributions based on rarefied datasets. The PERMANOVAs were calculated on normalised datasets by permuting 10,000 times with Site as a blocking factor, using Jaccard distances for 18S-V1V2 and COI, and Bray-Curtis distances for 18S-V4 and 16S-V4V5. Significant p values are in bold. For pairwise comparisons, DNA 10g comprises all DNA 10g processing methods, and significance codes are p<0.001: ‘***’; p<0.01: ‘**’; p<0.05: ‘*’.

| LOCUS | Cluster richness |  |  | Community differentiation |  |  |  |
| --- | --- | --- | --- | --- | --- | --- | --- |
|  | Chi-square | p-value | significant pairwise comparisons |  | R<sup>2</sup> | p-value | significant pairwise comparisons |
| **18S-V1V2** |  |  |  |  |  |  |  |
| Molecular processing | 50.3 | < **0.001** | DNA 2g < RNA 2g ***<br>DNA 10g > DNA 2g *** | Molecular processing | 0.06 | < **0.001** | DNA 2g / RNA 2g ***<br>DNA 10g / DNA 2g ***<br>DNA 10g / RNA 2g *** |
| Site (random effect) | 16.2 | < **0.001** |  | Site | 0.23 | < **0.001** |  |
|  |  |  |  | Molecular processing x Site | 0.19 | 0.16 |  |
| **COI** |  |  |  |  |  |  |  |
| Molecular processing | 57.3 | < **0.001** | DNA 2g > RNA 2g **<br>DNA 10g > DNA 2g *<br>DNA 10g > RNA 2g *** | Molecular processing | 0.09 | < **0.001** | DNA 2g / RNA 2g **<br>DNA 10g / DNA 2g *<br>DNA 10g / RNA 2g ** |
| Site (random effect) | 2.2 | 0.14 |  | Site | 0.20 | < **0.001** |  |
|  |  |  |  | Molecular processing x Site | 0.17 | **0.0013** |  |
| **18S-V4** |  |  |  |  |  |  |  |
| Molecular processing | 38.3 | < **0.001** | DNA 2g < RNA 2g ***<br>DNA 10g > DNA 2g ** | Molecular processing | 0.08 | < **0.001** | DNA 2g / RNA 2g **<br>DNA 10g / DNA 2g **<br>DNA 10g / RNA 2g ** |
| Site (random effect) | 15.9 | < **0.001** |  | Site | 0.35 | < **0.001** |  |
|  |  |  |  | Molecular processing x Site | 0.20 | < **0.001** |  |
| **16S-V4V5** |  |  |  |  |  |  |  |
| Molecular processing | 55.0 | < **0.001** | DNA 2g < RNA 2g ***<br>DNA 10g < RNA 2g *** | Molecular processing | 0.06 | < **0.001** | DNA 2g / RNA 2g ***<br>DNA 10g / DNA 2g **<br>DNA 10g / RNA 2g *** |
| Site (random effect) | 3.4 | 0.07 |  | Site | 0.57 | < **0.001** |  |
|  |  |  |  | Molecular processing x Site | 0.14 | < **0.001** |  |

**Figure 1.**
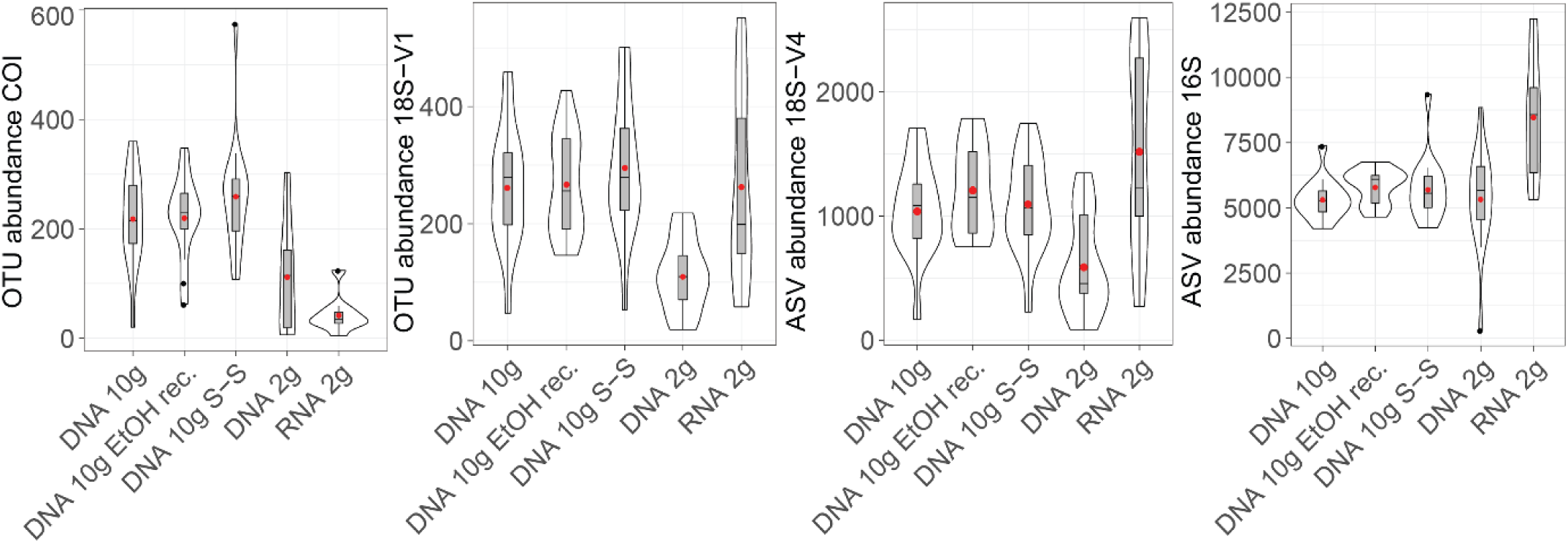
Violin plot showing detected numbers of metazoan OTUs (18S-V1, COI), micro-eukrayote (18S-V4) and prokaryote (16S-V4V5) ASVs recovered by the five molecular processing methods evaluated in this study. Cluster abundances were calculated on rarefied datasets. Boxplots show medians with interquartile ranges. Red dots indicate mean values.

Molecular processing designed to remove small DNA fragments (i.e. size-selection of DNA to remove fragments < 1,000 bp and ethanol reconcentration) did not significantly affect recovered cluster numbers obtained from eDNA extracted from 10 g of sediment, for any of the loci investigated (Fig. 1, Table 1, Tukey’s HSD multiple comparisons tests, p>0.9).

Extracts based on 2 g of sediment resulted in more variability, reflected by greater standard errors in mean recovered cluster numbers (15-26% of the mean for eukaryotes, 7-9% for prokaryotes) than in DNA extracts based on 10 g of sediment (8-11% for eukaryotes, 3-6% for prokaryotes).

DNA extracted from 2 g of sediment recovered significantly less eukaryotic clusters than extracts based on 10 g (Fig. 1, Table 1), a trend consistent across most taxa (Fig. 2). DNA-2g extracts recovered an average of 110±16 18S-V1 and 113±27 COI metazoan OTUs per sample, compared to 264±26 (18S-V1) and 222±23 (COI) in the DNA-10g extracts. Similarly, DNA-10g extracts recovered on average 1,117±100 protistan 18S-V4 ASVs per sample, compared to 595±109 detected in DNA from the 2 g kit. With 18S-V1, DNA extracted with the 2 g kit failed to detect three phyla (Brachiopoda, Loricifera, and Placozoa) that were present in both the RNA and the DNA-10g datasets. Contrastingly to eukaryotes, all DNA methods, whether based on 2 g or 10 g of sediment, resulted in comparable prokaryote ASV numbers detected (Figs. 1-2, Table 1, p>0.8), ranging from 5,330 ±199 to 5,810 ±170 per sample on average.

**Figure 2.**
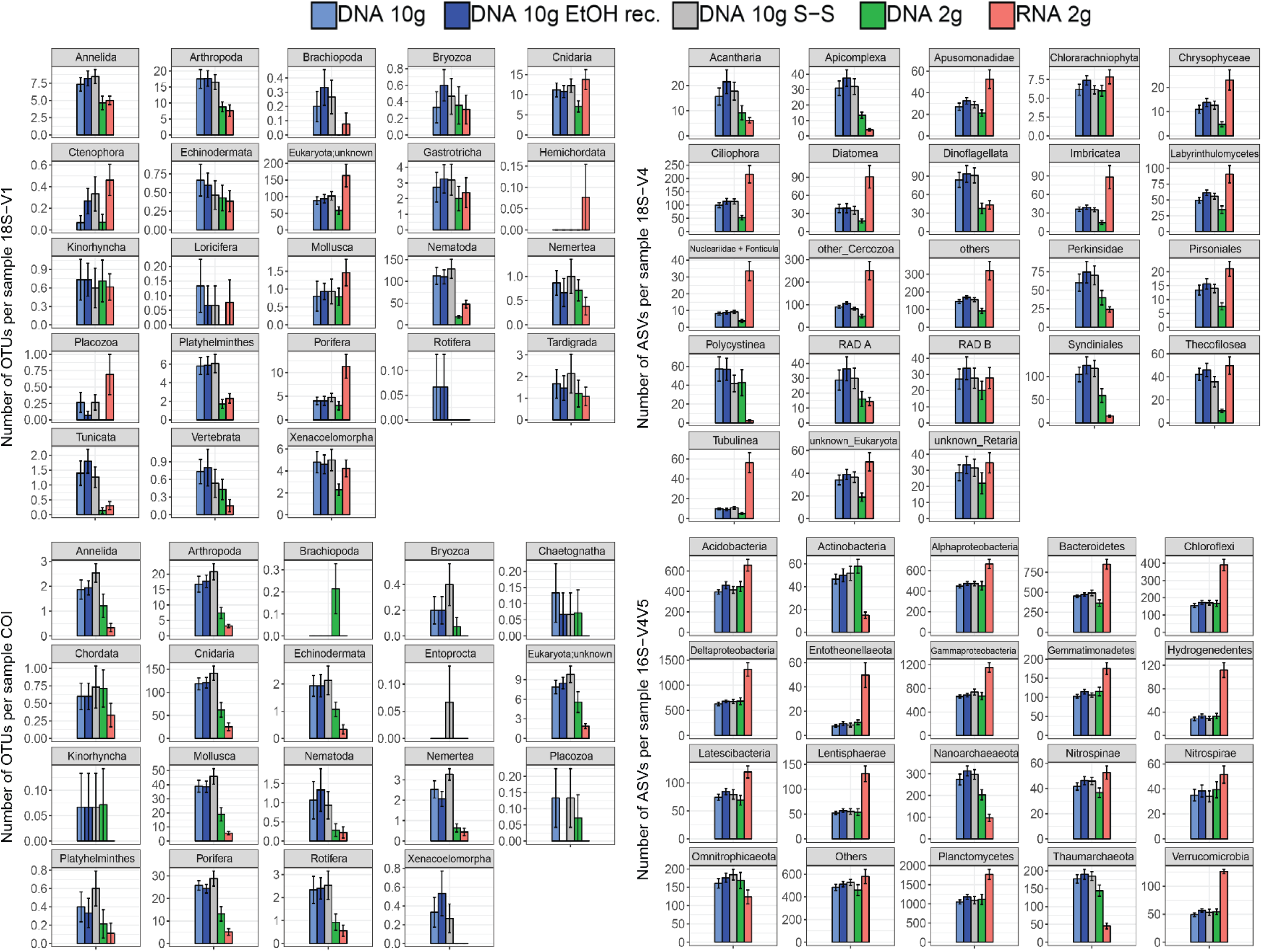
Mean number of metazoan OTUs (18S-V1, COI), protist ASVs (18S-V4), and prokaryote ASVs (16S-V4V5) per sample for each of the five processing methods. Cluster numbers were calculated on the rarefied datasets. Error bars represent standard errors.

The joint RNA/DNA extracts shared 15% (COI) to 25% (18S-V1) of metazoan OTUs, 14% of protistan 18S-V4 ASVs, and 25% of prokaryotic 16S ASVs (Fig. S 3). With COI, most unique OTUs were present in DNA extracts (74%), and RNA detected significantly less metazoan OTUs than co-extracted DNA (mean of 44±12 vs. 113±27 per sample), a trend observed in all detected metazoan phyla (Fig. 2). Contrastingly, with ribosomal loci, most clusters were unique to RNA (56% for 18S-V1, 63% for 18S-V4, 45% for 16S, Fig. S 3), which recovered significantly more clusters than co-extracted DNA (Fig. 1, Table 1). For prokaryotes, RNA extracts even detected significantly more ASVs than DNA extracts based on 10 g of sediment (Table 1, Fig. 1), a pattern observed in most prokaryotic clades, except for the Actinobacteria, Nanoarchaeaeota, Omnitrophicaeota, and the Thaumarchaeota (Fig. 2). For 18S-V4 and 18S-V1, RNA detected a cluster richness comparable to DNA-10 g extracts (Tukey’s HSD multiple comparisons tests, p>0.16), yet, average cluster numbers per sample were higher in RNA than in DNA-10g extracts in numerous groups (Fig. 2).

### 3.3. Effect of molecular processing methods on beta-diversity patterns

PERMANOVA showed that, although site was the main source of variation among samples (accounting for 20 to 57% of variability), significant differences existed among molecular methods in terms of community structure for all loci investigated over and above any variation due to site (Table 1). Pairwise comparisons indicated no significant effect of small DNA fragment removal on revealed community composition (Table 1), and high and significant correlations in Mantel tests (*r*: 0.92-1.0, p=0.001) confirmed the minor effect of size-selection and ethanol reconcentration. Based on these results, the size-selected and ethanol-reconcentrated DNA data were removed from further analyses, and community structures of the DNA-10g extracts were compared with those derived from co-extracted DNA/RNA using the 2g kit.

Pairwise comparisons showed significant differences in community structures between RNA and DNA for all markers analysed (Table 1). Ordinations, confirmed the predominant effect of site as the first two PCoA axes mostly resolved spatial effects (Figs S 4), but also revealed that communities detected by RNA differed from those detected by DNA (co-extracted DNA and DNA-10g), the level of differentiation varying among sites (Fig. 3).

**Figure 3.**
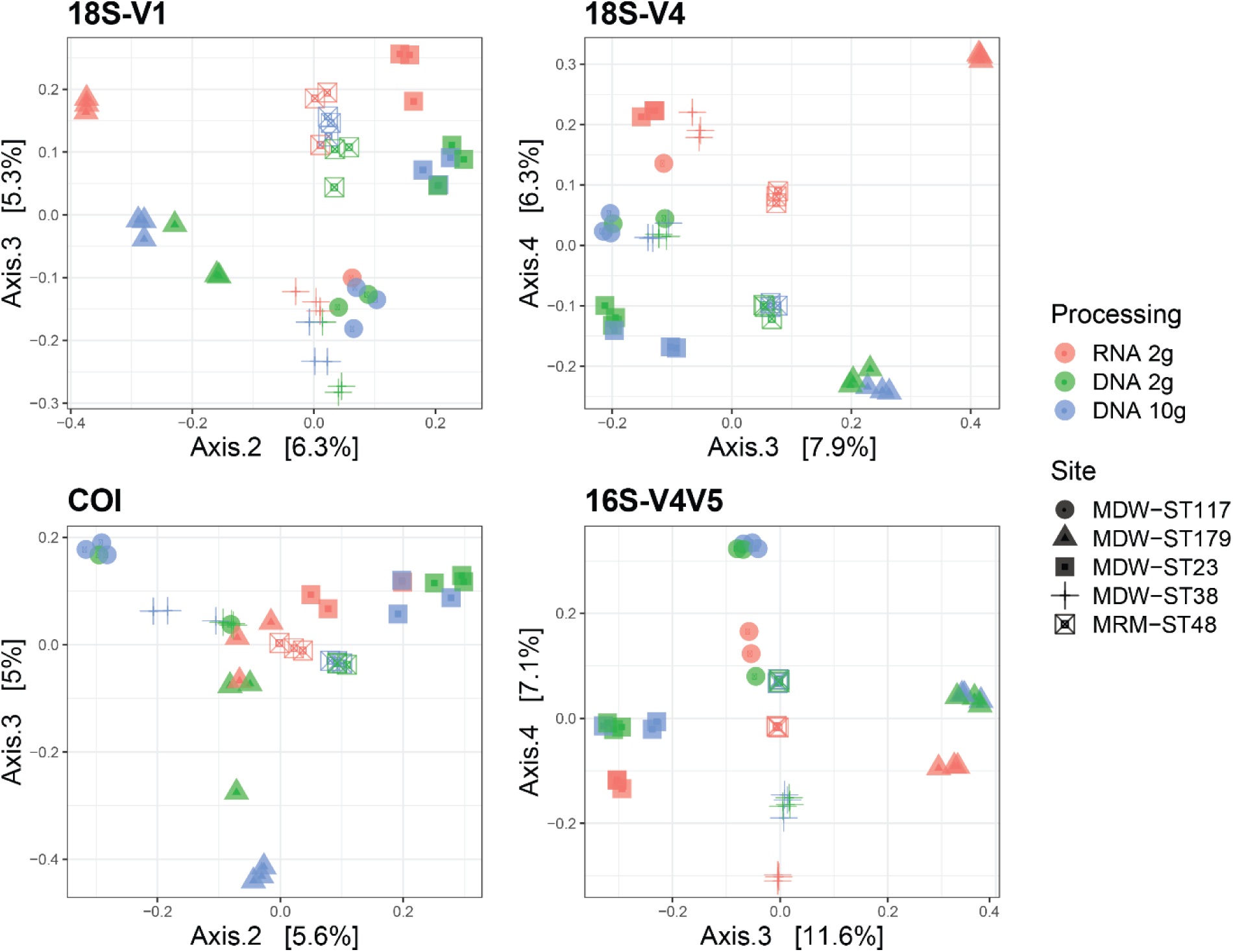
PCoA ordinations showing community differences between RNA and DNA molecular processing methods, using either DNA/RNA extracted jointly from 2 g of sediment or DNA extracted from 10g of sediment in five deep-sea sites using four barcode markers targeting metazoans (18S-V1, COI), micro-eukaryotes (18S-V4), and prokaryotes (16S-V4V5). PCoAs were calculated using Jaccard dissimilarities for metazoans and Bray-Curtis dissimilarities for unicellular organisms.

Pairwise comparisons also indicated significant differences in community structure between DNA extracts from the 2 g and 10 g kits (Table 1), possibly due to higher variability among replicate cores in the DNA-2g method as seen in ordinations (Fig. 3).

### 3.4. Extraction kit *vs* nature of nucleic acid

PERMANOVA of the dataset containing DNA-10g, DNA-2 g, and RNA-2g extracts confirmed that site was the predominant effect, explaining ∼20% of variation for metazoans, 33% of variation for micro-eukaryotes, and 54% of variation for prokaryotes. The analysis also indicated that the differences observed between processing methods were predominantly due to the type of nucleic acid rather than the kit used for extraction. Nucleic acid nature (DNA *vs* RNA) led to significant differences among assemblages for all loci, while DNA extraction kit resulted in significant differences only for 18S-V1 and 18S-V4 (Table S 6).

This supported observations in relative taxonomic compositions, which were more similar between samples based on DNA (Fig. 4), a pattern consistent across cores within each site (Fig. S 5). Expectedly, when looking at read numbers, resolved taxonomic structures were also more similar among DNA-based methods (Fig. S 6). Comparing read and cluster abundances revealed that relative taxonomic compositions based on read numbers (Fig. S 6) were comparable to those based on cluster numbers (Fig. 4) for micro-eukaryotes and prokaryotes, and confirmed that this was not the case for metazoans.

**Figure 4.**
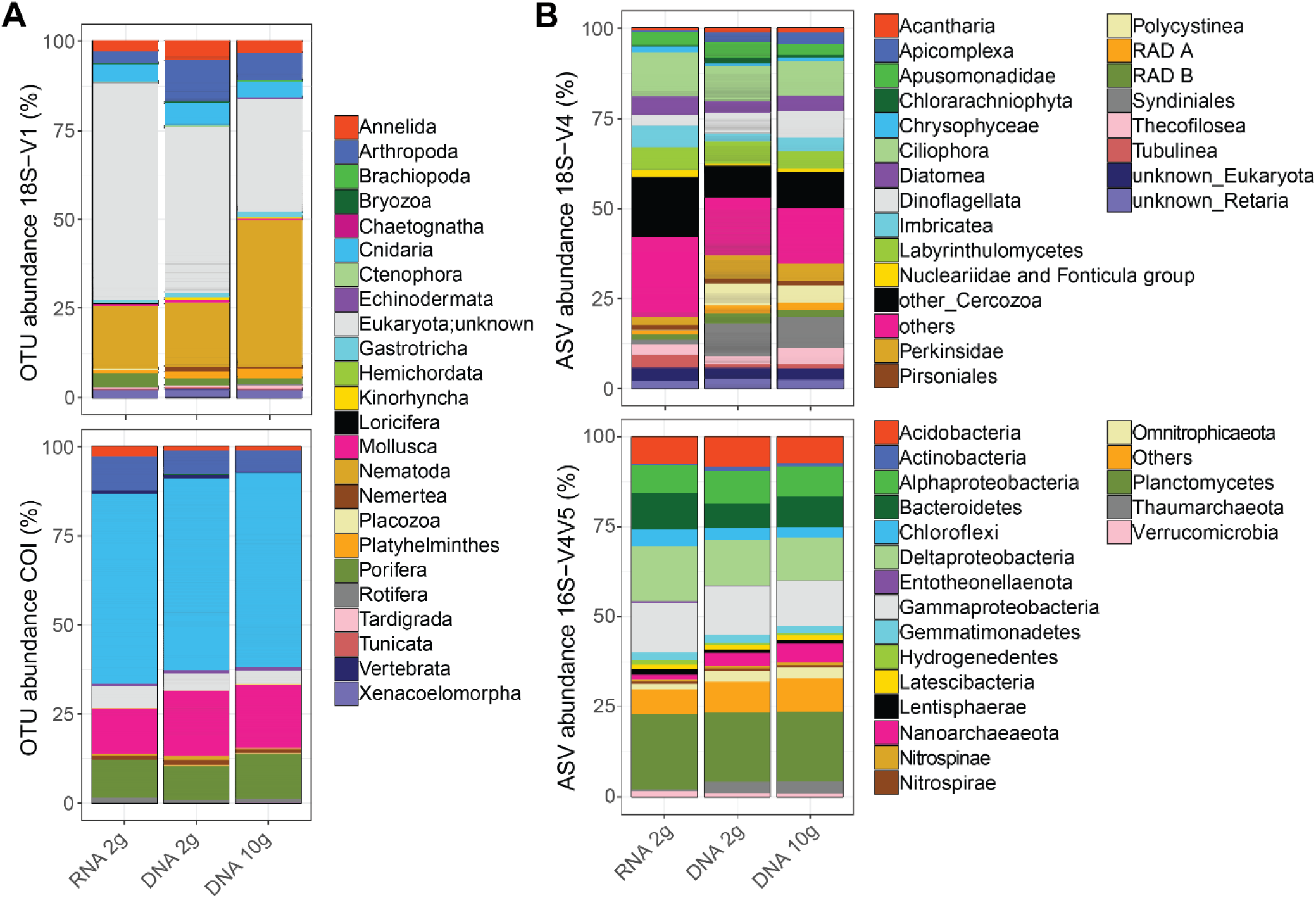
Patterns of relative cluster abundance resolved by metabarcoding results in five deep-sea sites by RNA and DNA molecular processing methods using either DNA/RNA extracted jointly from 2 g of sediment or DNA extracted from 10g of sediment, using four barcode markers targeting metazoans (**A**: 18S-V1, COI), micro-eukaryotes (**B**: 18S-V4), and prokaryotes (**B**: 16S-V4V5).

## 4. Discussion

The aim of this study was to evaluate different molecular methods in order to select the most appropriate eDNA metabarcoding protocol to inventory contemporary deep-sea communities, with the lowest possible bias due to aDNA.

Using RNA rather than DNA to inventory contemporaneous communities has been suggested as a means of avoiding the bias due to long-term persistence of DNA in marine sediments. Indeed, RNA is only produced by living organisms and is thought to quickly degrade when released in the environment, due to spontaneous hydrolysis and the abundance of RNases (Torti et al., 2015). Expectedly, in our COI dataset, RNA resulted in less OTUs (Fig. 1) and detected less phyla (Fig. 2) than co-extracted DNA. Contrastingly, for ribosomal loci, RNA detected higher cluster numbers than co-extracted DNA (Fig. 1), resulting in more clusters per sample for most of the taxonomic groups detected (Fig. 2). In these joint datasets, 45-63% of clusters were unique to RNA (Fig. S 2). These unique clusters were not singleton clusters as only up to 2.2% of them had less than three reads, even if 5-28% had less than ten reads (data not shown). Although proportions vary strongly among investigations, other studies using ribosomal loci have also reported increased recovery of OTUs in RNA datasets, and considerable amounts of unshared OTUs between joint RNA and DNA data (Guardiola et al., 2016; Laroche et al., 2017 and references therein).

This difference observed here between COI and ribosomal loci is likely related to the nature of the targeted RNA molecule. The rapid hydrolysis of RNA mostly applies to random coils (like mRNA), while helical conformations (including most types of RNA, such as ribosomal RNA, transfer RNA, viral genomic RNA, or ribozymes) are less prone to hydrolysis by water molecules (Torti et al., 2015). The degradation of rRNA is thus likely to be much slower than that of mRNA, which, combined with decreased digestion by RNases due to adsorption onto sediment particles (Torti et al., 2015), makes long-term persistence of rRNA possible, and observed in sediments and even in fossils (Cristescu, 2019; Orsi et al., 2013). Moreover, rRNA may generate excessive numbers of molecular clusters due to increased intraspecific polymorphism caused by stronger selection pressure on secondary structure rather than on rDNA sequences. Finally, the great abundance of RNA over DNA in living organisms (e.g. 20.5% vs 3.1% in *E. coli*) may also favour its persistence in the environment. This is especially true for rRNA, which is represented in a cell’s RNA pool as many times as there are ribosomes, while only being present in a few copies (10-150) in the genome (Torti et al., 2015).

While RNA has been reported as an effective way to depict the active community compartment (Baldrian et al., 2012; Lejzerowicz, Voltsky, et al., 2013; Pawlowski, Esling, Lejzerowicz, Cedhagen, & Wilding, 2014), variation in activity levels between taxonomic groups as well as differences in life histories, life strategies, and non-growth activities may confound this interpretation and generate taxonomic bias (Blazewicz, Barnard, Daly, & Firestone, 2013). Instead, DNA/RNA ratios might reflect different genomic architectures (variation in rDNA copy number) among taxonomic groups, rather than different relative activities (Massana et al., 2015). Thus, environmental RNA data need to be interpreted with caution, as some molecular clusters could be overrepresented due to increased cellular activities (Pochon, Zaiko, Fletcher, Laroche, & Wood, 2017). This could explain the higher cluster numbers detected for ribosomal loci with eRNA compared to eDNA for several taxa (Fig. 2).

Moreover, many of the unique RNA OTUs may be artefacts from the reverse transcription of RNA to cDNA, a process known to generate errors that are difficult to measure and detect in bioinformatic analyses (Laroche et al., 2017), but highlighted by the greater amounts of chimeras detected in RNA extracts with ribosomal loci. This overestimation of RNA-based data will affect non-clustered data more than clustered datasets, in line with the results observed here for microbial ASVs and metazoan OTUs.

In terms of beta diversity patterns, although RNA and DNA detected significantly different communities (Table 1, Fig. 3), DNA and RNA samples resolved similar spatial configurations, with samples clustering by site (Fig. 3). This is consistent with Guardiola et al. (2016), who also reported similar patterns of ecological differentiation between DNA and RNA in deep-sea sites although both datasets resolved different communities. However, spatial variation was more pronounced with DNA samples for eukaryotes, which is congruent with Laroche et al. (2017), who suggested that eDNA may be more reliable for assessing differences in community composition.

Thus, due to its suspected persistence in the environment, and the unknown but potentially additional sources of bias suspected here, using eRNA for metabarcoding of deep-sea sediments does not seem to effectively address the problem of aDNA, and even less so for ribosomal loci. Other studies suggested that a more efficient way to deal with aDNA may be to use joint RNA and DNA datasets, and trim for shared OTUs (Laroche et al., 2017; Pochon et al., 2017). This is however particularly stringent (given the low shared OTU proportions observed in this and other studies), and may result in a substantial number of false negatives. With COI, while mRNA may be more effectively targeting living organisms, the approach remains confronted with the taxonomic bias mentioned above, combined with higher *in vitro* lability of mRNA making it more challenging to work with (highlighted by the increased failure of RNA extracts in this study, Table S 5).

Removing small DNA fragments via size-selection (removing fragments < 1,000 bp) or ethanol reconcentration of DNA did not affect recovered cluster numbers in any of the biological compartments investigated (Fig. 1). The methods also did not result in any significant difference in community structures (Table 1), suggesting that small, likely ancient, extracellular DNA fragments have a negligible impact on biodiversity inventories produced through eDNA metabarcoding. This finding is in line with results from the deep-sea (Guardiola et al., 2016; Ramírez, Jørgensen, Zhao, & D’Hondt, 2018) and various other habitats (Lennon et al., 2018), which showed no evidence that spatial patterns were blurred by “dead” DNA persistence, and suggested a minimal effect of extracellular DNA on estimates of taxonomic and phylogenetic diversity.

None of the methods evaluated in the present study remove DNA not enclosed in *living* cells (e.g. DNA in organelles, DNA from dead cells…). It is still unclear how long DNA can remain intracellular after cell death or within organelles, and future research quantifying the rate at which “dead” intracellular DNA becomes extracellular and degraded will be valuable to estimate the potential bias of archived intracellular DNA in eDNA metabarcoding inventories of extant communities. However, there is increasing evidence that DNA from non-living cells is mostly contemporary (Lennon et al. 2018). This ability to detect extant taxa that were not present in the sample at the time of collection highlights the capacity of eDNA metabarcoding to detect local presence of organisms even from their remains or excretions, and even with a small amount of environmental material.

It remains to be elucidated whether more cost and time effective extraction protocols specifically targeting extracellular DNA offer similar ecological resolution as total DNA kits. This is suggested to be the case for terrestrials soils (Taberlet, Prud’Homme, et al., 2012; Zinger et al., 2016), although authors have highlighted that conclusions from these studies should be interpreted with caution as results might be influenced by actively released and ancient DNA (Nagler et al. 2018). The only available study testing this in the deep-sea showed that richness patterns were strikingly different in several metazoan phyla between extracellular DNA and total DNA. The authors suggested this to be the result of activity bias: sponges and cnidarians were overrepresented in the extracellular DNA pool because they continuously expel DNA, while nematodes were underrepresented as their cuticles shield DNA (Guardiola et al., 2016). As this comparison was performed on samples collected in two consecutive years, differences observed may partly result from temporal variation. However, another study of shallow and mesobenthic macroinvertebrates showed that targeting solely the extracellular eDNA compartment of marine sediments led to the detection of more than 100 taxa less than bulk metabarcoding or morphology, suggesting that extracellular DNA may not be adequate for marine sediments (Aylagas, Borja, Irigoien, & Rodríguez-Ezpeleta, 2016).

Larger amounts of sediment (≥10g) allowed detecting significantly more eukaryotic clusters. This was not true for prokaryotes, for which both 2 g and 10 g of sediment detected similar numbers of ASVs (Table 1, Fig. 1). It may be suggested that in joint the RNA/DNA kit, DNA elution occurring after RNA elution induces partial DNA loss. However, such effect would be expected to equally affect eu- and prokaryotes, which was not the case here, supporting the fact that quantity of starting material significantly affects results for eukaryotes. The importance of adjusting the amount of starting material to the biological compartment investigated has already been documented (Creer et al., 2016; Dopheide, Xie, Buckley, Drummond, & Newcomb, 2019), and this study confirms that while 2-4 g of deep-sea sediment may be enough to capture prokaryote diversity, microbial eukaryotes and metazoans are more effectively surveyed with larger sediment volumes.

Finally, the 2 g protocols were generally associated to results with higher variability among replicate cores for all loci investigated (Fig. 1, Fig. 3). This variability increases confidence intervals, reduces statistical power, and increases the risk of not identifying differences among communities, and thus impacts in EIA studies (Type II errors). Small-scale (cm to metres) patchiness has often been reported in the deep-sea (Grassle & Maciolek, 1992; Lejzerowicz, Esling, & Pawlowski, 2014; Smith & Snelgrove, 2002). While technical (PCR) replicates allow increasing taxon detection probability (decrease false positives), this within-site variability can only be mitigated by collecting more biological replicates per sampling station, and using a sufficiently high amount of starting material to extract nucleic acids.

## ACKNOWLEDGEMENTS

This work is part of the “*Pourquoi Pas les Abysses?*” project funded by Ifremer, and the project eDNAbyss (AP2016 -228) funded by France Génomique (ANR-10-INBS-09) and Genoscope-CEA. This work also received funding from the European Union’s Horizon 2020 research and innovation programme under grant the agreement No 678760 (ATLAS). This output reflects only the author’s view and the European Union cannot be held responsible for any use that may be made of the information contained therein. We wish to thank Laure Quintric, Patrick Durand, and Caroline Belser for bioinformatic support, as well as Jan Pawlowski and Eva Ramirez Llodra for useful advice on this work. We also wish to express our gratitude to the crew, participants, and mission chiefs of the MarMine cruise (Eva Ramirez Llodra, project 247626/O30 (NRC-BI) and the MEDWAVES cruise supported by the ATLAS project and the Spanish Ministry of Economy, Industry and Competitivity (Covadonga Orejas and all the crew from the Sarmiento de Gamboa), as well as to all the people who helped collecting samples (Perregrino Cambeiro, Juancho Movilla, Maria Rakka, Joana Boavida, Anna Addamo).

## AUTHOR CONTRIBUTIONS

MIB, CLH and SAH designed the study, MIB, JP, and CLH carried out the laboratory work. MIB and BT performed the bioinformatic analyses. MIB, BT, and NH performed the statistical analyses. MIB and SAH wrote the manuscript. All authors contributed to the final manuscript.

## DATA ACCESSIBILITY

The data for this work can be accessed in the European Nucleotide Archive (ENA) database (Study accession number will be given upon manuscript acceptance). The dataset, including sequences, databases, as well as raw and refined ASV/OTU tables, have been deposited on ftp://ftp.ifremer.fr/ifremer/dataref/bioinfo/merlin/abyss/abyss-molecular-comparisons/. Bioinformatic scripts and config files are available on Gitlab (https://gitlab.ifremer.fr/abyss-project/).

## Notes

ftp://ftp.ifremer.fr/ifremer/dataref/bioinfo/merlin/abyss/abyss-molecular-comparisons/

https://gitlab.ifremer.fr/abyss-project

## REFERENCES

Aylagas, E., Borja, Á., Irigoien, X., & Rodríguez-Ezpeleta, N. (2016). Benchmarking DNA Metabarcoding for Biodiversity-Based Monitoring and Assessment. Frontiers in Marine Science, 3. https://doi.org/10.3389/fmars.2016.00096

Baldrian, P., Kolaiřík, M., Štursová, M., Kopecký, J., Valášková, V., Větrovský, T., … Voříšková, J. (2012). Active and total microbial communities in forest soil are largely different and highly stratified during decomposition. ISME Journal, 6(2), 248–258. https://doi.org/10.1038/ismej.2011.95

Baselga, A., & Orme, C. D. L. (2012). betapart : an R package for the study of beta diversity. Methods in Ecology and Evolution, 3(5), 808–812. https://doi.org/10.1111/j.2041-210X.2012.00224.x

Blazewicz, S. J., Barnard, R. L., Daly, R. A., & Firestone, M. K. (2013). Evaluating rRNA as an indicator of microbial activity in environmental communities: Limitations and uses. ISME Journal, 7(11), 2061–2068. https://doi.org/10.1038/ismej.2013.102

Boere, A. C., Rijpstra, W. I. C., De Lange, G. J., Sinninghe Damsté, J. S., & Coolen, M. J. L. (2011). Preservation potential of ancient plankton DNA in Pleistocene marine sediments. Geobiology, 9(5), 377–393. https://doi.org/10.1111/j.1472-4669.2011.00290.x

Brandt, M. I., Trouche, B., Quintric, L., Wincker, P., Poulain, J., & Arnaud-Haond, S. (2019). A flexible pipeline combining bioinformatic correction tools for prokaryotic and eukaryotic metabarcoding. BioRxiv, 717355. https://doi.org/10.1101/717355

Callahan, B. J., McMurdie, P. J., Rosen, M. J., Han, A. W., Johnson, A. J. A., & Holmes, S. P. (2016). DADA2: High-resolution sample inference from Illumina amplicon data. Nature Methods, 13(7), 581–583. https://doi.org/10.1038/nmeth.3869

Carugati, L., Corinaldesi, C., Dell’Anno, A., & Danovaro, R. (2015). Metagenetic tools for the census of marine meiofaunal biodiversity: An overview. Marine Genomics, 24, 11–20. https://doi.org/10.1016/j.margen.2015.04.010

Coolen, M. J. L., Orsi, W. D., Balkema, C., Quince, C., Harris, K., Sylva, S. P., … Giosan, L. (2013). Evolution of the plankton paleome in the Black Sea from the Deglacial to Anthropocene. Proceedings of the National Academy of Sciences of the United States of America, 110(21), 8609–8614. https://doi.org/10.1073/pnas.1219283110

Corinaldesi, C., Beolchini, F., & Dell’Anno, A. (2008). Damage and degradation rates of extracellular DNA in marine sediments: Implications for the preservation of gene sequences. Molecular Ecology, 17(17), 3939–3951. https://doi.org/10.1111/j.1365-294X.2008.03880.x

Corinaldesi, Cinzia, Tangherlini, M., Manea, E., & Dell’Anno, A. (2018). Extracellular DNA as a genetic recorder of microbial diversity in benthic deep-sea ecosystems. Scientific Reports, 8(1), 1839. https://doi.org/10.1038/s41598-018-20302-7

Creer, S., Deiner, K., Frey, S., Porazinska, D., Taberlet, P., Thomas, W. K., … Bik, H. M. (2016). The ecologist’s field guide to sequence-based identification of biodiversity. Methods in Ecology and Evolution, 7(9), 1008–1018. https://doi.org/10.1111/2041-210X.12574

Cristescu, M. E. (2019). Can Environmental RNA Revolutionize Biodiversity Science? Trends in Ecology and Evolution. https://doi.org/10.1016/j.tree.2019.05.003

Davis, N. M., Proctor, D. M., Holmes, S. P., Relman, D. A., & Callahan, B. J. (2018). Simple statistical identification and removal of contaminant sequences in marker-gene and metagenomics data. Microbiome, 6(1), 226. https://doi.org/10.1186/s40168-018-0605-2

Dell’Anno, A., & Danovaro, R. (2005). Ecology: Extracellular DNA plays a key role in deep-sea ecosystem functioning. Science, 309(5744), 2179. https://doi.org/10.1126/science.1117475

Dopheide, A., Xie, D., Buckley, T. R., Drummond, A. J., & Newcomb, R. D. (2019). Impacts of DNA extraction and PCR on DNA metabarcoding estimates of soil biodiversity. Methods in Ecology and Evolution, 10(1), 120–133. https://doi.org/10.1111/2041-210X.13086

Escudié, F., Auer, L., Bernard, M., Mariadassou, M., Cauquil, L., Vidal, K., … Berger, B. (2018). FROGS: Find, Rapidly, OTUs with Galaxy Solution. Bioinformatics, 34(8), 1287–1294. https://doi.org/10.1093/bioinformatics/btx791

Frøslev, T. G., Kjøller, R., Bruun, H. H., Ejrnæs, R., Brunbjerg, A. K., Pietroni, C., & Hansen, A. J. (2017). Algorithm for post-clustering curation of DNA amplicon data yields reliable biodiversity estimates. Nature Communications, 8(1). https://doi.org/10.1038/s41467-017-01312-x

Grassle, J. F., & Maciolek, N. J. (1992). Deep-sea species richness: regional and local diversity estimates from quantitative bottom samples. American Naturalist, 139(2), 313–341. https://doi.org/10.1086/285329

Guardiola, M., Wangensteen, O. S., Taberlet, P., Coissac, E., Uriz, M. J., & Turon, X. (2016). Spatio-temporal monitoring of deep-sea communities using metabarcoding of sediment DNA and RNA. PeerJ, 4, e2807. https://doi.org/10.7717/peerj.2807

Laroche, O., Wood, S. A., Tremblay, L. A., Ellis, J. I., Lear, G., & Pochon, X. (2018). A cross-taxa study using environmental DNA/RNA metabarcoding to measure biological impacts of offshore oil and gas drilling and production operations. Marine Pollution Bulletin, 127, 97–107. https://doi.org/10.1016/j.marpolbul.2017.11.042

Laroche, O., Wood, S. A., Tremblay, L. A., Lear, G., Ellis, J. I., & Pochon, X. (2017). Metabarcoding monitoring analysis: The pros and cons of using co-extracted environmental DNA and RNA data to assess offshore oil production impacts on benthic communities. PeerJ, 2017(5), e3347. https://doi.org/10.7717/peerj.3347

Lejzerowicz, F., Esling, P., Majewski, W., Szczuciński, W., Decelle, J., Obadia, C., … Pawlowski, J. (2013). Ancient DNA complements microfossil record in deep-sea subsurface sediments. Biology Letters, 9(4), 20130283. https://doi.org/10.1098/rsbl.2013.0283

Lejzerowicz, F., Esling, P., & Pawlowski, J. (2014). Patchiness of deep-sea benthic Foraminifera across the Southern Ocean: Insights from high-throughput DNA sequencing. Deep-Sea Research Part II: Topical Studies in Oceanography, 108, 17–26. https://doi.org/10.1016/j.dsr2.2014.07.018

Lejzerowicz, F., Voltsky, I., & Pawlowski, J. (2013). Identifying active foraminifera in the Sea of Japan using metatranscriptomic approach. Deep-Sea Research Part II: Topical Studies in Oceanography, 86–87, 214–220. https://doi.org/10.1016/j.dsr2.2012.08.008

Lennon, J. T., Muscarella, M. E., Placella, S. A., & Lehmkuhl, B. K. (2018). How, when, and where relic DNA affects microbial diversity. MBio, 9(3), e00637–18. https://doi.org/10.1128/mBio.00637-18

Leray, M., Yang, J. Y., Meyer, C. P., Mills, S. C., Agudelo, N., Ranwez, V., … Machida, R. J. (2013). A new versatile primer set targeting a short fragment of the mitochondrial COI region for metabarcoding metazoan diversity: application for characterizing coral reef fish gut contents. Front Zool, 10, 34. https://doi.org/10.1186/1742-9994-10-34

Machida, R. J., Leray, M., Ho, S. L., & Knowlton, N. (2017). Data Descriptor: Metazoan mitochondrial gene sequence reference datasets for taxonomic assignment of environmental samples. Scientific Data, 4. https://doi.org/10.1038/sdata.2017.27

Massana, R. R., Gobet, A., Audic, S., Bass, D., Bittner, L., Boutte, C., … de Vargas, C. (2015). Marine protist diversity in European coastal waters and sediments as revealed by high-throughput sequencing. Environmental Microbiology, 17(10), 4035–4049. https://doi.org/10.1111/1462-2920.12955

McMurdie, P. J., & Holmes, S. (2013). Phyloseq: An R Package for Reproducible Interactive Analysis and Graphics of Microbiome Census Data. PLoS ONE, 8(4), e61217. https://doi.org/10.1371/journal.pone.0061217

Nagler, M., Insam, H., Pietramellara, G., & Ascher-Jenull, J. (2018). Extracellular DNA in natural environments: features, relevance and applications. Applied Microbiology and Biotechnology, 102(15), 6343–6356. https://doi.org/10.1007/s00253-018-9120-4

Oksanen, J., Blanchet, Guillaume F. Friendly, M., Kindt, R., Legendre, P., McGlinn, D., Minchin, R. P., … Wagner, H. (2018). vegan: Community Ecology Package. Retrieved from https://cran.r-project.org/package=vegan

Orsi, W., Biddle, J. F., & Edgcomb, V. (2013). Deep Sequencing of Subseafloor Eukaryotic rRNA Reveals Active Fungi across Marine Subsurface Provinces. PLoS ONE, 8(2), e56335. https://doi.org/10.1371/journal.pone.0056335

Parada, A. E., Needham, D. M., & Fuhrman, J. A. (2016). Every base matters: assessing small subunit rRNA primers for marine microbiomes with mock communities, time series and global field samples. Environ Microbiol, 18(5), 1403–1414. https://doi.org/10.1111/1462-2920.13023

Pawlowski, J. W., Esling, P., Lejzerowicz, F., Cedhagen, T., & Wilding, T. A. (2014). Environmental monitoring through protist next-generation sequencing metabarcoding: assessing the impact of fish farming on benthic foraminifera communities. Mol Ecol Resour, 14(6), 1129–1140. https://doi.org/10.1111/1755-0998.12261

Pochon, X., Zaiko, A., Fletcher, L. M., Laroche, O., & Wood, S. A. (2017). Wanted dead or alive? Using metabarcoding of environmental DNA and RNA to distinguish living assemblages for biosecurity applications. PLoS ONE, 12(11), e0187636. https://doi.org/10.1371/journal.pone.0187636

Quast, C., Pruesse, E., Yilmaz, P., Gerken, J., Schweer, T., Yarza, P., … Glöckner, F. O. (2012). The SILVA ribosomal RNA gene database project: improved data processing and web-based tools. Nucleic Acids Research, 41(D1), D590–D596. https://doi.org/10.1093/nar/gks1219

R Core Team. (2018). R: A language and environment for statistical computing. R Foundation for Statistical Computing, Vienna, Austria.

Ramirez-Llodra, E., Tyler, P. A., Baker, M. C., Bergstad, O. A., Clark, M. R., Escobar, E., … van Dover, C. L. (2011). Man and the last great wilderness: Human impact on the deep sea. PLoS ONE, 6(7), e22588. https://doi.org/10.1371/journal.pone.0022588

Ramírez, G. A., Jørgensen, S. L., Zhao, R., & D’Hondt, S. (2018). Minimal Influence of Extracellular DNA on Molecular Surveys of Marine Sedimentary Communities. Frontiers in Microbiology, 9, 2969. https://doi.org/10.3389/fmicb.2018.02969

Sinniger, F., Pawlowski, J. W., Harii, S., Gooday, A. J., Yamamoto, H., Chevaldonné, P., … Creer, S. (2016). Worldwide analysis of sedimentary DNA reveals major gaps in taxonomic knowledge of deep-sea benthos. Frontiers in Marine Science, 3(June), 92. https://doi.org/10.3389/FMARS.2016.00092

Smith, C., & Snelgrove, P. (2002). A Riot of Species in An Environmental Calm: The Paradox of the Species-Rich Deep-Sea Floor (pp. 311–342). CRC Press. https://doi.org/10.1201/9780203180594.ch6

Stoeck, T., Bass, D., Nebel, M., Christen, R., Jones, M. D. M., Breiner, H. W., & Richards, T. A. (2010). Multiple marker parallel tag environmental DNA sequencing reveals a highly complex eukaryotic community in marine anoxic water. Molecular Ecology, 19(SUPPL. 1), 21–31. https://doi.org/10.1111/j.1365-294X.2009.04480.x

Taberlet, P., Coissac, E., Hajibabaei, M., & Rieseberg, L. H. (2012). Environmental DNA. Molecular Ecology, 21(8), 1789–1793. https://doi.org/10.1111/j.1365-294X.2012.05542.x

Taberlet, P., Prud’Homme, S. M., Campione, E., Roy, J., Miquel, C., Shehzad, W., … Coissac, E. (2012). Soil sampling and isolation of extracellular DNA from large amount of starting material suitable for metabarcoding studies. Molecular Ecology, 21(8), 1816–1820. https://doi.org/10.1111/j.1365-294X.2011.05317.x

Torti, A., Lever, M. A., & Jørgensen, B. B. (2015). Origin, dynamics, and implications of extracellular DNA pools in marine sediments. Marine Genomics. https://doi.org/10.1016/j.margen.2015.08.007

Wangensteen, O. S., & Turon, X. (2016). Metabarcoding Techniques for Assessing Biodiversity of Marine Animal Forests. In S. Rossi, L. Bramanti, A. Gori, & C. Orejas Saco del Valle (Eds.), Marine Animal Forests (pp. 1–29). Cham: Springer International Publishing. https://doi.org/10.1007/978-3-319-17001-5_53-1

Zinger, L., Chave, J., Coissac, E., Iribar, A., Louisanna, E., Manzi, S., … Taberlet, P. (2016). Extracellular DNA extraction is a fast, cheap and reliable alternative for multi-taxa surveys based on soil DNA. Soil Biology and Biochemistry, 96, 16–19. https://doi.org/10.1016/j.soilbio.2016.01.008

