## Supplementary material for "An assessment of environmental metabarcoding protocols aiming at favouring contemporary biodiversity in inventories of deep-sea communities"

### SUPPLEMENTAL INFORMATION

### 1 Supplemental materials and methods

#### 1.1 Collection of samples

Table S 1. Sampling sites and their GPS locations and associated habitats. MEDWAVES: MEDiterranean outflow WAter and Vulnerable EcosystemS.

| Site | Cruise | Depth (m) | Latitude | Longitude | Habitat | Region |
|---|---|---|---|---|---|---|
| MDW-ST179; Seco de los Olivos | MEDWAVES | 729 | 36.4808 | -2.8945 | Seamount | Western Mediterranean |
| MDW-ST23; Gazul | MEDWAVES | 470 | 36.5605 | -6.9498 | Mud volcano | Gibraltar Strait |
| MDW-ST38; Ormonde | MEDWAVES | 1,920 | 36.8442 | -11.3025 | Seamount | North Atlantic |
| MDW-ST117; Formigas | MEDWAVES | 1,325 | 37.34 | -24.7552 | Seamount | North Atlantic |
| MRM-ST48; Mohn’s Treasure | MarMine | 2,826 | 73.4598 | 7.2184 | Hydrothermal vent | Arctic |

#### 1.2 PCR amplification

Table S 2. Primers used in this study, targeting metazoans with the COI and 18S-V1V2 loci, micro-eukaryotes with the 18S-V4 barcode, and prokaryotes with the 16S-V4V5 marker.

| Locus | Target and *specificity* | Primer forward Primer reverse | Short name | Sequence (5'-3') | Amplicon size (bp) | Reference |
|---|---|---|---|---|---|---|
| **COI** | Eukaryotes | mlCOIintF | COI-F | GGWACWGGWTGAACWGTWTAYCCYCC | 313 | Leray et al., 2013 |
| | *pref. metazoans* | jgHCO2198 | COI-R | TAIACYTCIGGRTGICCRAARAAYCA | | |
| **18S-V1V2** | Eukaryotes | SSUF04 | 18S-V1F | GCTTGTCTCAAAGATTAAGCC | 330-390 | Sinniger et al., 2016 |
| | *pref. metazoans* | SSURmod | 18S-V1-R | CCTGCTGCCTTCCTTRGA | | |
| **18S-V4** | Eukaryotes | V4F (TAReukFWD1) | 18S-V4-F | CCAGCASCYGCGGTAATTCC | 350-410 | Stoeck et al., 2010 |
| | *all* | V4R (TAReukREV3) | 18S-V4-R | ACTTTCGTTCTTGATYRA | | |
| **16S-V4V5** | Prokaryotes | 515f | 16S-F | GTGYCAGCMGCCGCGGTAA | 350-390 | Parada et al., 2016 |
| | *Pref. Eubacteria* | 926r | 16S-R | CCGYCAATTYMTTTRAGTTT | | |

##### *1.2.1 Eukaryotic 18S-V1V2 rRNA gene amplicon generation*

Eukaryotic 18S-V1V2 barcodes were generated using the SSUF04 (5'-GCTTGTCTCAAAGATTAAGCC-3') and SSUR22*mod* (5'-CCTGCTGCCTTCCTTRGA-3') primers (Sinniger et al., 2016) and the *Phusion* High Fidelity PCR Master Mix with GC buffer (ThermoFisher Scientific, Waltham, MA, USA). The PCR reactions (25 µL final volume) contained 2.5 ng or less of DNA template with 0.4 µM concentration of each primer, 3% of DMSO, and 1X Phusion Master Mix.

PCR amplifications (98 °C for 30 s; 25 cycles of 10 s at 98 °C, 30 s at 45 °C, 30 s at 72 °C; and 72 °C for 10 min) of all samples were carried out in triplicate in order to smooth the intra-sample variance while obtaining sufficient amounts of amplicons for Illumina sequencing. Amplicon triplicates were pooled and PCR products were purified using 1X AMPure XP beads (Beckman Coulter, Brea, CA, USA) cleanup. Aliquots of purified amplicons were run on an Agilent Bioanalyzer using the DNA High Sensitivity LabChip kit (Agilent Technologies, Santa Clara, CA, USA) to check their lengths, and quantified with a Qubit fluorometer (Invitrogen, Carlsbad, CA, USA).

##### *1.2.2 Eukaryotic 18S-V4 rRNA gene amplicon generation*

Eukaryotic 18S-V4 barcodes were generated using the TAReukF1 (5'-CCAGCASCYGCGGTAATTCC-3') and TAReukR (5'-ACTTTCGTTCTTGATYRA-3') primers (Stoeck et al., 2010). Triplicate PCR reactions were prepared as described above, but amplification was performed by a nested PCR with the first annealing temperature being 53°C for 10 cycles, followed by 48°C for 15 cycles. After PCR product cleanup using 1X AMPure XP beads, amplicon lengths and amounts were checked as described above.

##### *1.2.3 Prokaryotic 16S-V4V5 rRNA gene amplicon generation*

Prokaryotic barcodes were generated using the 515F-Y (5′-GTGYCAGCMGCCGCGGTAA-3′) and 926R (5′- CCGYCAATTYMTTTRAGTTT-3′**)**

primers (Parada, Needham, & Fuhrman, 2016). Triplicate PCR reactions were prepared as described above for 18S-V1V2, but annealing temperature was at 53 °C. After PCR product cleanup using 1X AMPure XP beads, amplicon lengths and amounts were checked as described above.

###### *1.2.4 Eukaryotic COI gene amplicon generation*

Metazoan COI barcodes were generated using the mlCOIintF 5'-GGWACWGGWTGAACWGTWTAYCCYCC-3'and jgHCO2198 5'-TAIACYTCIGGRTGICCRAARAAYCA-3' primers (Leray et al. 2013). The PCR reactions (20 µL final volume) contained 2.5 ng or less of total DNA template with 0.5 µM final concentration of each primer, 3% of DMSO, 0.175 mM final concentration of dNTPs, and 1X Advantage 2 Polymerase Mix (Takara Bio, Kusatsu, Japan). Nested PCR amplifications were carried out in triplicates and consisted of an initial denaturation at 95 °C for 10 min, and 16 cycles of 10 s at 95°C, 30 s at 62 °C (−1°C per cycle), 60 s at 68 °C followed by 15 cycles of 95 °C for 10 s, 30 s at 46°C, 68 °C for 60 s, and a final extension of 68 °C for 7 min.

#### 1.3 High Throughput Sequencing

###### *1.3.1 Amplicon library preparation*

One hundred ng of amplicons were directly end-repaired, A-tailed and ligated to Illumina adapters on a Biomek FX Laboratory Automation Workstation (Beckman Coulter, Brea, CA, USA). Library amplification was performed using a Kapa Hifi HotStart NGS library Amplification kit (Kapa Biosystems, Wilmington, MA, USA) with the same cycling conditions applied for all metagenomic libraries and purified using 1X AMPure XP beads.

###### *1.3.2 Sequencing library quality control*

Libraries were quantified by Quant-iT dsDNA HS assay kits using a Fluoroskan Ascent microplate fluorometer (Thermo Fisher Scientific, Waltham, MA, USA) and then by qPCR with the KAPA Library Quantification Kit for Illumina Libraries (Kapa Biosystems, Wilmington, MA, USA) on a MxPro instrument (Agilent Technologies, Santa Clara, CA, USA). Library profiles were assessed using a high-throughput microfluidic capillary electrophoresis system (LabChip GX, Perkin Elmer, Waltham, MA, USA).

###### 1.1.1 Sequencing procedures

Library concentrations were normalized to 10 nM by addition of 10 mM Tris-Cl (pH 8.5) and applied to cluster generation according to the Illumina Cbot User Guide (Part # 15006165). Amplicon libraries are characterized by low diversity sequences at the beginning of the reads due to the presence of the primer sequence. Low-diversity libraries can interfere in correct cluster identification, resulting in a drastic loss of data output. Therefore, loading concentrations of libraries were decreased (8–9 pM instead of 12–14 pM for standard libraries) and PhiX DNA spike-in was increased (20% instead of 1%) in order to minimize the impacts on the run quality.

Libraries were sequenced on HiSeq2500 (System User Guide Part # 15035786) instruments (Illumina, San Diego, CA, USA) in 250 base pairs paired-end mode.

#### 1.4 Bioinformatic analyses

Table S 3. ABYSS metabarcoding pipeline.

| Process | Software | Script(s) and command(s) |
|---|---|---|
| Raw reads preprocessing for ligation data | Abyss-preprocessing: separate forward and reverse reads in each run, and re-pair reads | extractR1R2.pbs using cutadapt v1.18 (-e 0.17 for 18S-V1 and 0.27 for COI , -O length of primer -1) and BBMAP Repair v38.22 |
| Read quality-filtering | Dada2 v.1.10 | filterAndTrim() in dada2main.R maxEE=2, maxN=0, truncQ=11, truncLen=220 (18S, 16S) or 200 (COI) |
| Read error learning | Dada2 v.1.10 | learnErrors() in dada2main.R nbases=1e8, multithread=TRUE, randomize=TRUE |
| Read dereplicating | Dada2 v.1.10 | derepFastq() in dada2main.R |
| Read correction | Dada2 v.1.10 | dada() in dada2main.R |
| Read merging | Dada2 v.1.10 | mergePairs() in dada2main.R minOverlap=12, maxMismatch=0 |
| Make sequence table and filter by length | Dada2 v.1.10 | makeSequenceTable() in dada2main.R seqtab[,nchar(colnames(seqtab)) %in% seq(lengthMin,lengthMax)] lengthMin= 330 (18S-V1), 300 (COI), 350 (18S-V4), 350 (16S) lengthMax= 390 (18S-V1), 326 (COI), 410 (18S-V4), 390 (16S) |
| Chimera removal | Dada2 v.1.10 | removeBimeraDenovo() in dada2main.R |
| Taxonomic assignment with RDP Classifier | Dada2 v.1.10 | assignTaxonomy () in dada2outputfiles.R minBoot=50, outputBootstraps=TRUE |
| Taxonomic assignment with BLAST+ | blastn (megablast) v.2.6.0 | blast.pbs -outfmt 11 -qcov_hsp_perc 80 -perc_identity 70 -max_hsps 1, -evalue 1e-5, then merge BLAST and RDP taxonomies using concat_blast_rdp_tax.pbs |
| Clustering (optional) | FROGS v.2.0.0 | clustering.py with d=4 for 18S-V1V2 and d=6 for COI, remova_chimera.py, affiliation_OTU_identite_couverture.py |
| Blank correction | Rscript | Data_refining.Rmd using packages decontam v.1.2.1 and phyloseq v.1.26.0 |
| Removal of unassigned and non-target clusters | | |
| Deletion of defective samples (< 10,000 target reads) | | |
| Tag-switching renormalisation | Rscript | owi_renormalize.R |
| LULU curation | LULU v.0.1 | lulu() in lulu_final.R minimum_ratio_type = "min", minimum_ratio = 1, minimum_match =84, minimum_relative_cooccurence =0.93 |

### 2 Supplemental tables

Table S 4. Tests for homogeneity of multivariate dispersions for the 4 genes studied. The tests for performed with 9999 permutations on Jaccard distances for 18S-V1 and COI, and on Bray-Curtis distances for 18S-V4 and 16S. Significant p values are in bold. For pairwise comparisons, DNA 10g comprises all DNA 10g processing methods, and significance codes are: p<0.001: '***'; p<0.01: '**'; p<0.05: '*'. In cases of significantly different dispersions, PERMANOVAs were performed on balanced datasets.

| LOCUS | *df* | SS | F-value | *p-value* | significant parwise comparisons |
|---|---|---|---|---|---|
| **18S-V1** | | | | | |
| Molecular processing | 4 | 0.00270 | 1.26 | 0.3 | NA |
| Residuals | 67 | 0.0361 | | | |
| **COI** | | | | | |
| Molecular processing | 4 | 0.00349 | 3.26 | **0.017** | RNA 2g / DNA 10g *<br>DNA 2g / DNA 10g S-S * |
| Residuals | 63 | 0.0169 | | | |
| **18S-V4** | | | | | |
| Molecular processing | 4 | 0.00222 | 0.78 | 0.54 | NA |
| Residuals | 64 | 0.0450 | | | |
| **16S-V4V5** | | | | | |
| Molecular processing | 4 | 0.00875 | 0.93 | 0.44 | NA |
| Residuals | 69 | 0.161 | | | |

Table S 5. Number of reads and clusters (ASVs for 18S-V4 and 16S, OTUs for 18S-V1 and COI) obtained at different analysis steps, depending on molecular processing. Data refining was performed in R, based on BLAST assignments obtained using the Silva v132 database for 18S and 16S loci, and on the MIDORI database for COI. Final number of target reads represent the number of target-taxa reads after data refining (abundance renormalisation for 18S and 16S loci, abundance renormalisation and LULU curation for COI). Final number of target clusters are the corresponding ASVs for 18S-V4 and 16S, and the corresponding OTUs for 18S-V1 and COI.

| Sample type | Number of samples | Raw reads | Quality-filtered reads | Merged reads | Length-filtered reads | Non chimeric reads | % reads retained | Number of clusters before refining | Number of samples after refining | Final number of target reads | Final number of target clusters |
|---|---|---|---|---|---|---|---|---|---|---|---|
| **LOCUS** | | | | | | | | | | | |
| **18S-V1V2** | | | | | | | | | | | |
| Control | 5 | 2,921,651 | 1,654,366 | 1,601,427 | 1,380,613 | 1,379,141 | 47 | 42,876 | 0 | 16,157,973 | 6,031 |
| DNA 10g | 15 | 12,969,202 | 9,978,581 | 9,141,929 | 8,577,414 | 8,463,482 | 65 | | 15 | | |
| DNA 10g EtOH rec. | 15 | 13,646,370 | 10,577,221 | 9,757,954 | 9,271,161 | 9,129,915 | 67 | | 15 | | |
| DNA 10g S-S | 15 | 11,735,871 | 8,938,926 | 7,990,574 | 7,403,206 | 7,328,224 | 62 | | 15 | | |
| DNA 2g | 14 | 8,476,073 | 6,402,605 | 5,840,025 | 5,215,391 | 5,168,422 | 61 | | 14 | | |
| Positive Control (Metazoa only) | 2 | 2,096,631 | 1,607,219 | 1,438,424 | 1,432,399 | 1,293,985 | 62 | | 2 | | |
| RNA 2g | 14 | 18,130,054 | 14,322,872 | 13,311,099 | 12,180,021 | 11,591,521 | 64 | | 13 | | |
| **COI** | | | | | | | | | | | |
| Control | 7 | 642,571 | 414,329 | 410,866 | 410,260 | 410,189 | 64 | 45,508 | 0 | 10,977,614 | 4,333 |
| DNA 10g | 15 | 13,804,664 | 11,871,147 | 11,645,233 | 10,515,311 | 10,437,446 | 76 | | 15 | | |
| DNA 10g EtOH rec. | 15 | 12,735,345 | 10,966,940 | 10,758,928 | 9,634,212 | 9,560,863 | 75 | | 15 | | |
| DNA 10g S-S | 15 | 13,172,416 | 11,357,075 | 11,141,979 | 10,019,802 | 9,948,428 | 76 | | 15 | | |
| DNA 2g | 14 | 10,992,972 | 8,962,857 | 8,748,635 | 7,478,953 | 7,439,814 | 68 | | 14 | | |
| Positive Control (Metazoa only) | 2 | 1,482,785 | 1,261,045 | 1,253,408 | 1,252,485 | 1,226,728 | 83 | | 2 | | |
| RNA 2g | 9 | 8,085,884 | 6,548,055 | 6,405,869 | 5,780,511 | 5,749,188 | 71 | | 9 | | |
| **18S-V4** | | | | | | | | | | | |
| Control | 5 | 38,028 | 1,088 | 1,005 | 786 | 786 | 2 | 65,832 | 0 | 8,654,710 | 40,868 |
| DNA 10g | 15 | 5,108,793 | 3,852,156 | 3,244,507 | 3,081,831 | 3,073,436 | 60 | | 15 | | |
| DNA 10g EtOH rec. | 13 | 4,812,187 | 3,622,684 | 3,014,540 | 2,884,756 | 2,876,930 | 60 | | 13 | | |
| DNA 10g S-S | 15 | 3,675,283 | 2,779,263 | 2,334,265 | 2,222,758 | 2,216,696 | 60 | | 15 | | |
| DNA 2g | 13 | 2,569,170 | 1,853,940 | 1,539,838 | 1,422,958 | 1,419,415 | 55 | | 13 | | |
| RNA 2g | 13 | 14,024,345 | 10,695,784 | 8,260,707 | 6,978,244 | 6,876,335 | 49 | | 13 | | |
| **16S-V4V5** | | | | | | | | | | | |
| Control | 5 | 1,100,024 | 815,505 | 692,998 | 687,737 | 686,244 | 62 | 148,797 | 0 | 21,740,351 | 138,478 |
| DNA 10g | 15 | 6,228,145 | 4,351,718 | 3,445,745 | 3,436,831 | 3,311,742 | 53 | | 15 | | |
| DNA 10g EtOH rec. | 15 | 7,400,388 | 5,167,853 | 4,033,571 | 4,024,943 | 3,861,002 | 52 | | 15 | | |
| DNA 10g S-S | 15 | 7,163,763 | 5,039,022 | 4,045,902 | 4,037,814 | 3,891,760 | 54 | | 15 | | |
| DNA 2g | 15 | 7,788,742 | 5,409,390 | 4,582,930 | 4,573,875 | 4,428,719 | 57 | | 15 | | |
| RNA 2g | 14 | 15,248,792 | 10,719,709 | 7,858,978 | 7,854,365 | 7,438,875 | 49 | | 14 | | |

Table S 6. PERMANOVAs of the DNA-10g/DNA-2g/RNA-2g datasets for the 4 studied genes. The PERMANOVAs were calculated on normalised datasets by permuting 10,000 times with Site as a blocking factor, using Jaccard distances for 18S-V1 and COI, and Bray-Curtis distances for 18S-V4 and 16S-V4V5. Significant p values are in bold.

| **LOCUS** | **18S-V1** | | **COI** | | **18S-V4** | | **16S-V4V5** | |
|---|---|---|---|---|---|---|---|---|
| | **R^2** | ***p-value*** | **R^2** | ***p-value*** | **R^2** | ***p-value*** | **R^2** | ***p-value*** |
| DNAvsRNA | 0.04 | < **0.001** | 0.04 | < **0.01** | 0.05 | < **0.001** | 0.05 | < **0.001** |
| Kit | 0.03 | < **0.001** | 0.038 | 0.09 | 0.027 | < **0.01** | 0.01 | 0.11 |
| Site | 0.23 | < **0.001** | 0.18 | < **0.001** | 0.33 | < **0.001** | 0.54 | < **0.001** |

### 3 Supplemental figures

#### 3.1 Alpha diversity between processing methods

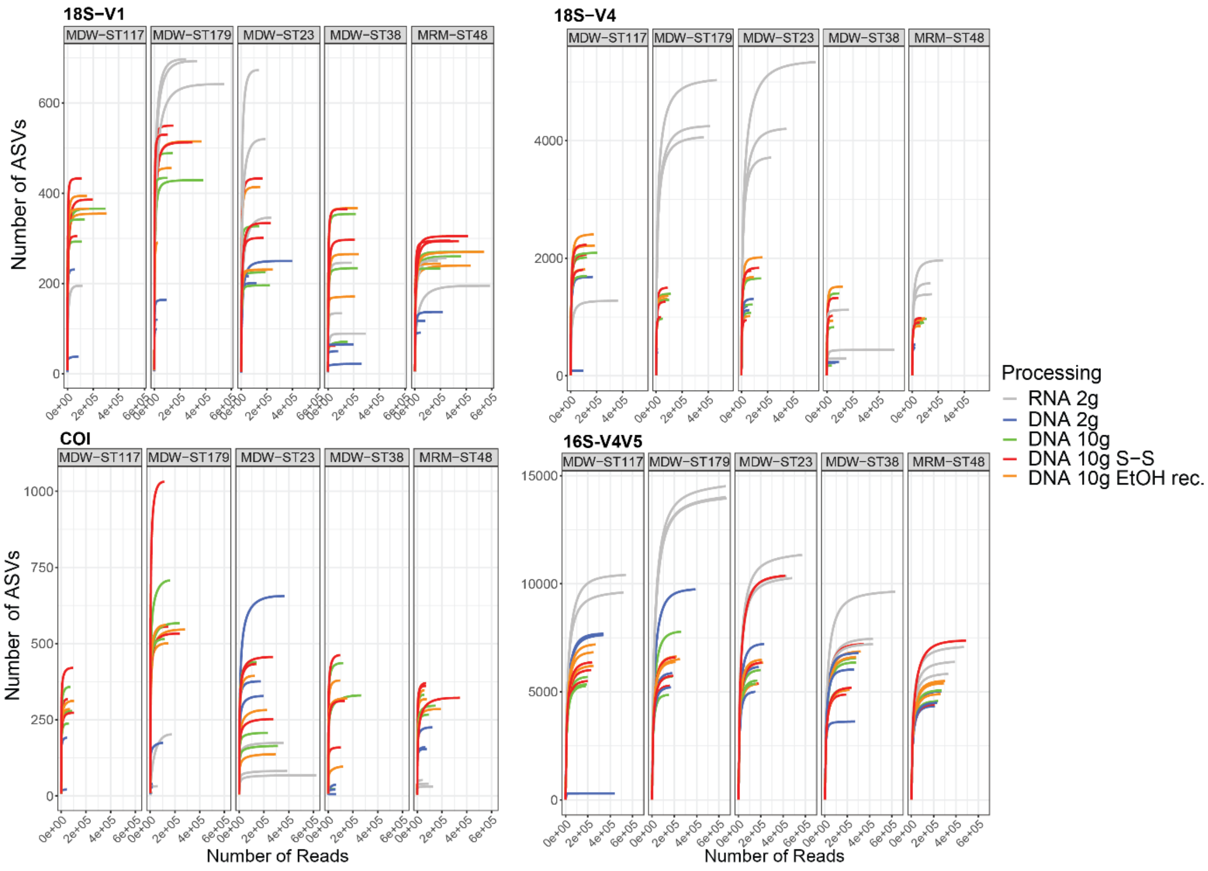


Figure S 1. Rarefaction curves in deep-sea sediment samples from 5 sampling sites, processed with 5 molecular methods for producing metabarcoding inventories of metazoans (18S-V1, COI), micro-eukaryotes (18S-V4), and prokaryotes (16S-V4V5), showing a plateau is reached in all samples.

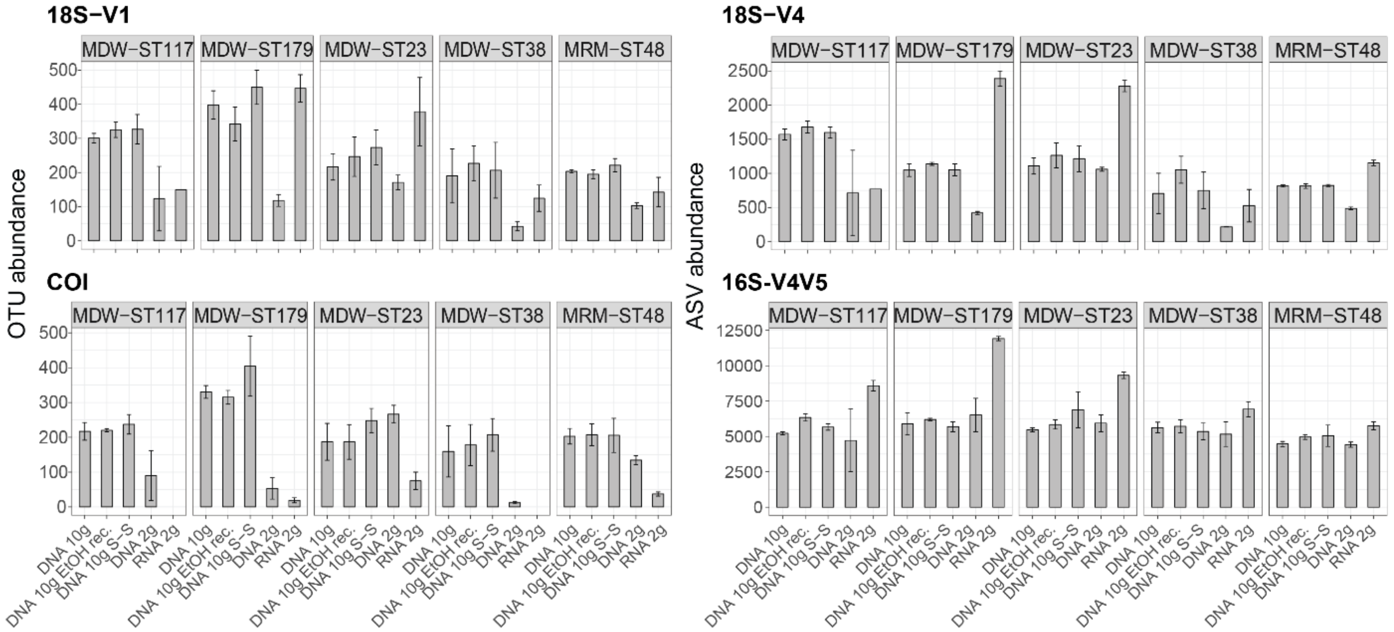


Figure S 2. Mean number of metazoan OTUs (18S-V1, COI), protist (18S-V4) and prokaryote ASVs (16S-V4V5) recovered by the five molecular processing methods evaluated in this study, in each of the five sampling sites. Cluster numbers were calculated on the rarefied datasets. Error bars represent standard errors.

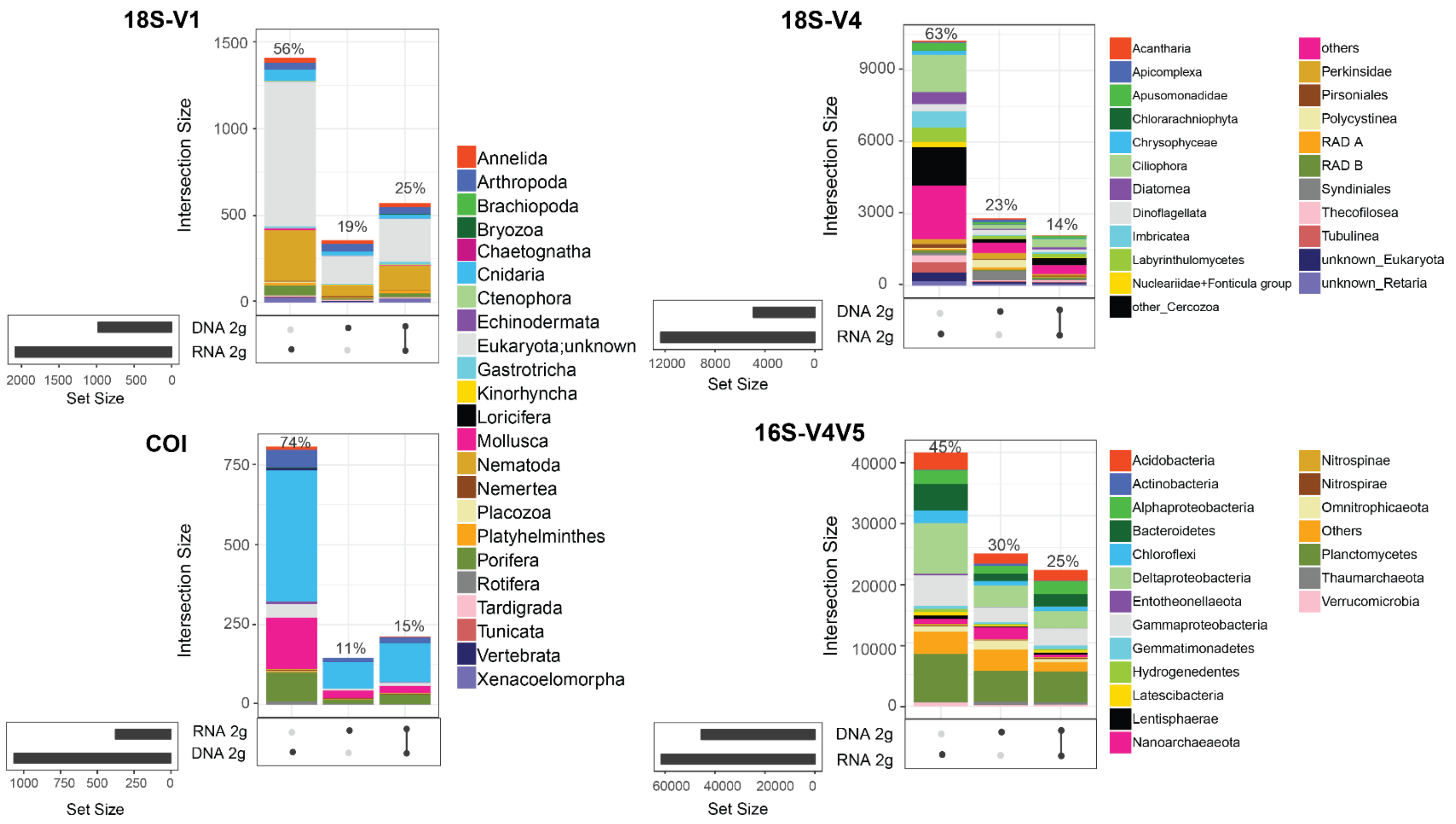


Figure S 3.Shared and unique metazoan OTUs (18S-V1, COI), protozoan ASVs (18S-V4), and prokaryote ASVs (16S-V4V5) among the joint DNA and RNA datasets. Numbers were calculated on the rarefied datasets.

#### 3.2 Effect of processing methods on beta-diversity patterns

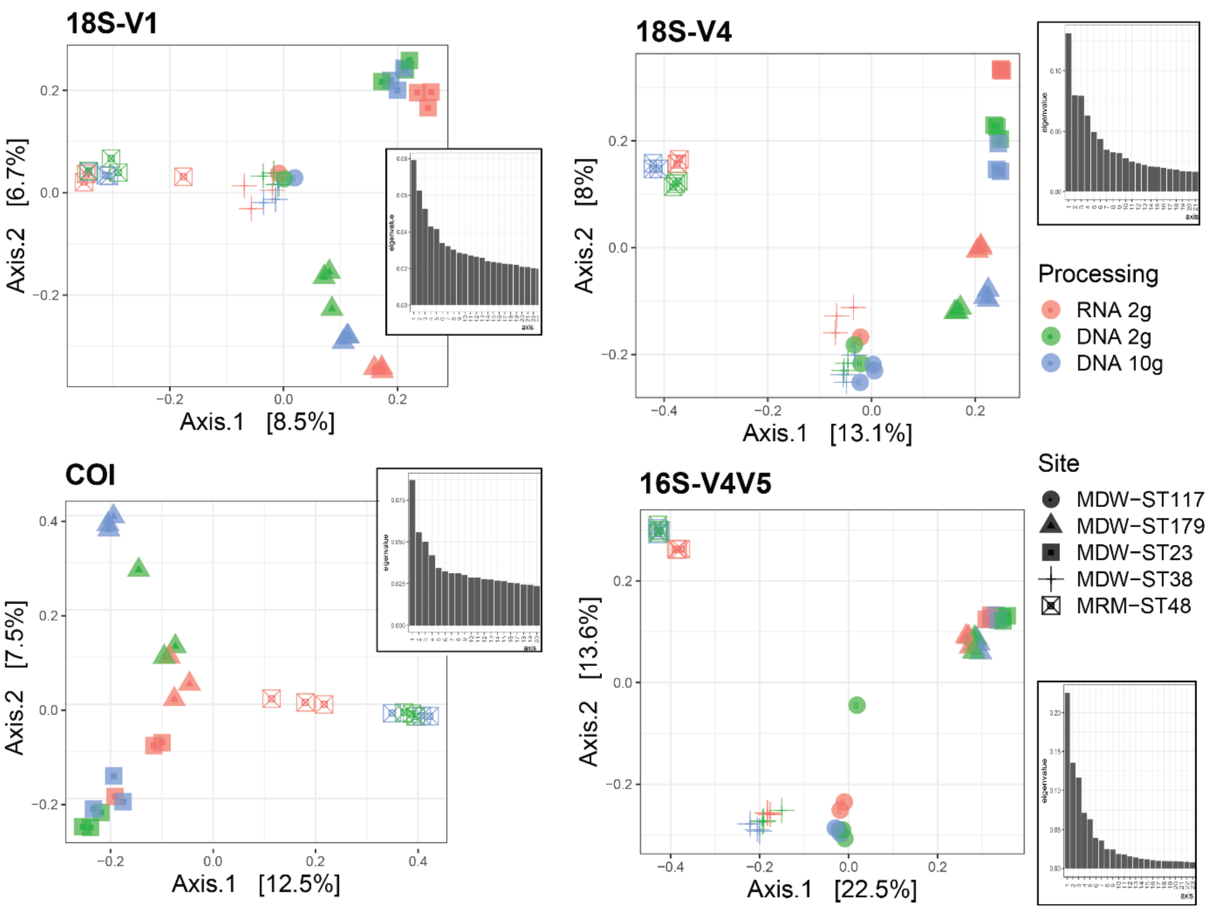


Figure S 4. Community differences between RNA and DNA molecular processing methods using either DNA/RNA extracted jointly from 2 g of sediment or DNA extracted from 10g of sediment in five deep-sea sites using four barcode markers targeting metazoans (18S-V1, COI), micro-eukaryotes (18S-V4), and prokaryotes (16S-V4V5). PCoAs were calculated using Jaccard dissimilarities for metazoans and Bray-Curtis dissimilarities for unicellular organisms. The first two axes of the PCoAs shown here capture the main source of variation, the site variation. Scree plots of each ordination are shown in inserts, indicating that variation due to processing method is captured by secondary axes.

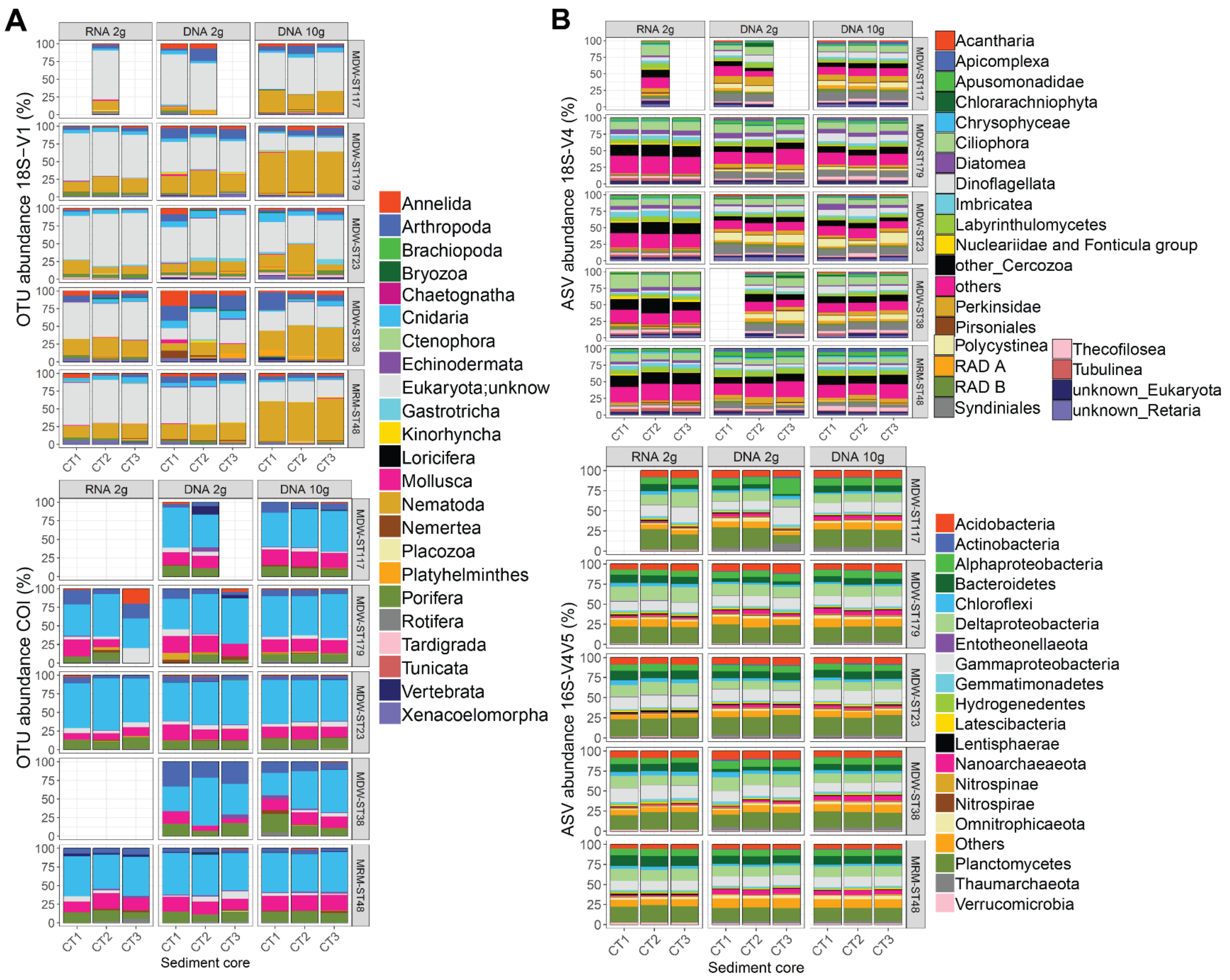


Figure S 5. Patterns of relative cluster abundance resolved by metabarcoding results in triplicate sediment cores from five deep-sea sites by RNA and DNA molecular processing methods using DNA/RNA extracted jointly from 2 g of sediment or DNA extracted from 10g of sediment, using four barcode markers targeting metazoans (**A**: 18S-V1, COI), micro-eukaryotes (**B**: 18S-V4), and prokaryotes (**B**: 16S-V4V5).

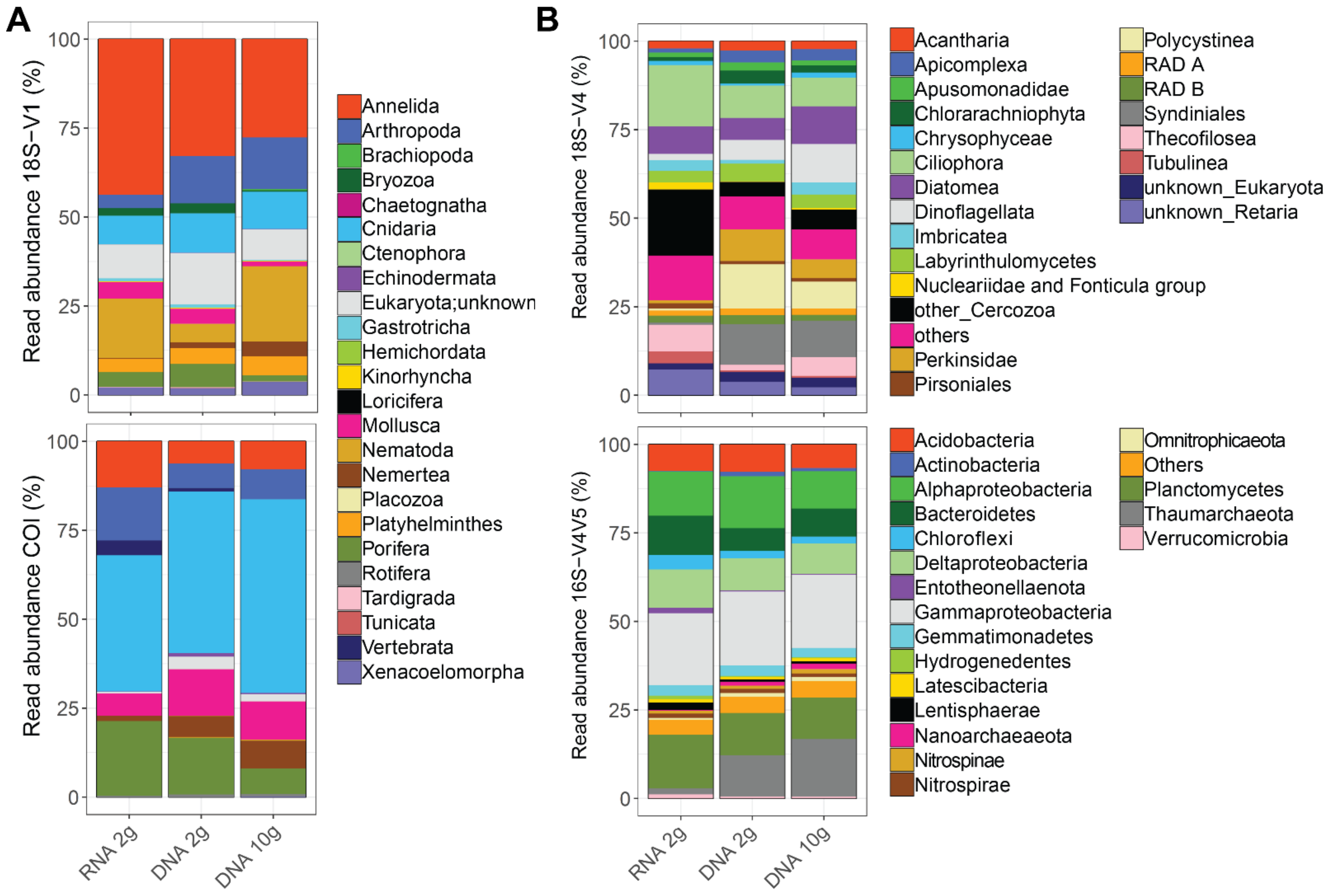


Figure S 6. Patterns of relative read abundance resolved by metabarcoding results in five deep-sea sites by RNA and DNA molecular processing methods using DNA/RNA extracted jointly from 2 g of sediment or DNA extracted from 10g of sediment, using four barcode markers targeting metazoans (**A**: 18S-V1, COI), micro-eukaryotes (**B**: 18S-V4), and prokaryotes (**B**: 16S-V4V5).
